# Microbiota of the pregnant mouse: characterization of the bacterial communities in the oral cavity, lung, intestine, and vagina through culture and DNA sequencing

**DOI:** 10.1101/2022.04.15.488507

**Authors:** Jonathan M. Greenberg, Roberto Romero, Andrew D. Winters, Jose Galaz, Valeria Garcia-Flores, Marcia Arenas-Hernandez, Jonathan Panzer, Zachary Shaffer, David J. Kracht, Nardhy Gomez-Lopez, Kevin R. Theis

**Author notes:** CORRESPONDING AUTHOR(S) Kevin R. Theis, Nardhy Gomez-Lopez.

## Abstract

Mice are frequently used as animal models for mechanistic studies of infection and obstetrical disease, yet characterization of the murine microbiota during pregnancy is lacking. The objective of this study was to therefore characterize the microbiotas of distinct body sites of the pregnant mouse that harbor microorganisms that could potentially invade the murine amniotic cavity leading to adverse pregnancy outcomes: vagina, oral cavity, intestine, and lung. The microbiotas of these body sites were characterized through anoxic, hypoxic, and oxic culture, as well as through 16S rRNA gene sequencing. With the exception of the vagina, the cultured microbiotas of each body site varied with atmosphere, with the greatest diversity in the cultured microbiota appearing under anoxic conditions. Only cultures of the vagina were able to recapitulate the microbiota observed from direct DNA sequencing of body site samples, primarily due to the dominance of two *Rodentibacter* strains. Identified as *R. pneumotropicus* and *R. heylii,* these isolates exhibited dominance patterns similar to those of *Lactobacillus crispatus* and *L. iners* in the human vagina. Whole genome sequencing of these *Rodentibacter* strains revealed shared genomic features, including the ability to degrade glycogen, an abundant polysaccharide in the vagina. In summary, we report body site specific microbiotas in the pregnant mouse with potential ecological parallels to those of humans. Importantly, our findings indicate that the vaginal microbiota of pregnant mice can be readily cultured, suggesting that mock vaginal microbiotas can be tractably generated and maintained for experimental manipulation in future mechanistic studies of host vaginal-microbiome interactions.

**IMPORTANCE:** Mice are widely utilized as animal models of obstetrical complications; however, the characterization of the murine microbiota has been neglected during pregnancy. Microorganisms from the vagina, oral cavity, intestine, and lung have been found in the intra-amniotic space, where their presence threatens the progression of gestation. Herein, we characterize the microbiotas of pregnant mice and establish the appropriateness of culture in capturing the microbiota at each site. The high relative abundance of *Rodentibacter* observed in the vagina is similar to that of *Lactobacillus* in humans, suggesting potential ecological parallels. Importantly, we report that the vaginal microbiota of the pregnant mouse can be readily cultured under hypoxic conditions, demonstrating that mock microbial communities can be utilized to test the potential ecological parallels between microbiotas in human and murine pregnancy, and to evaluate the relevance of the structure of these microbiotas for adverse pregnancy outcomes, especially intra-amniotic infection and spontaneous preterm birth.

## INTRODUCTION

Ethical and practical limitations on experimentation with humans are barriers to fully understanding the role of the microbiome in human health and disease. To overcome these limitations, researchers often perform experiments with *in vitro* cell culture models or *in vivo* animal models, presuming that these models accurately reflect host-microbiome dynamics in humans. In particular, the laboratory mouse is widely used for *in vivo* experimentation evaluating microbial causes of disease (1, 2). The mouse model has several benefits. First, of the available mammalian models, mice are relatively inexpensive to maintain and easy to manipulate experimentally. They can be housed in controlled environments, including environments that are germ-free, thereby reducing the impact of potential confounding variables on the microbiota and experimental outcomes related to health and disease. However, the mouse model is often used without consideration of the differences between the microbiotas of mice and humans or the potential differential impacts of the microbiota on health and disease in the two species (1, 3, 4). Specifically, experimental mouse studies often include the introduction of a human-specific microorganism into the mouse’s microbiota or the transplantation of an entire body site-specific human microbiota into the mouse. A limitation of these studies is a lack of knowledge of a mouse’s typical microbiota, making experimentally induced changes in the microbiota hard to interpret. This is further exacerbated by studies operating under the assumption that the human microbiota can be equivalently recreated within the mouse model or that interactions between the human microbiota and a mouse are the same as those between the human microbiota and a human (2). However, if parallels between the microbial ecology of human and mouse body site-specific microbiotas can be identified, and if mouse microbiotas can be tractably constituted through culture, manipulated in a targeted way, and reintroduced to the mouse, then focusing on the mouse microbiota in investigations of mouse models of health and disease may be as or more fruitful than focusing on the human microbiota.

The intestinal microbiota of the mouse has been intensively studied and characterized (5–20). However, the microbiotas of the mouse oral cavity, lung, and vagina have received much less attention **(Supplemental Tables 1-3)**, and only a select few studies have simultaneously characterized the microbiotas of multiple body sites in the mouse (5, 18, 21). This gap in knowledge is particularly apparent in studies of the mouse microbiota during pregnancy. The mouse has been widely used to investigate pregnancy complications, including perinatal infection and spontaneous preterm birth (22–38); yet, aside from the intestinal microbiota (15, 17, 39), the microbiota of the mouse in the context of pregnancy has been largely overlooked **(Table 1)**. Given the widely reported associations between the human vaginal microbiota and pregnancy complications, such as intra-amniotic infection (40–43) and spontaneous preterm birth (36, 44–58), the baseline vaginal microbiota in the pregnant mouse should be thoroughly investigated and characterized. This is critical because the human vaginal microbiota is unique – humans are the only mammal known to have vaginal microbiotas that are often dominated by just a single bacterial species (i.e., one of four *Lactobacillus* spp.: principally *L. crispatus* or *L. iners,* and secondarily *L. gasseri* or *L. jensenii*) (59–61), and these microbiotas have been characterized into readily distinguishable vaginal community state types (CSTs) (62). These *Lactobacillus-* dominant CSTs (CSTs I-III, and V) are generally perceived as being conducive to reproductive health. Conversely, the relationship between human reproductive health and the non- *Lactobacillus-*dominant, and thus species rich and diverse, CST IV is more ambiguous (45, 55- 57, 63–65). This disparity in health outcome potentially related to the structure of the vaginal microbiota is highlighted by the observation that women who do not have a *Lactobacillus*-dominant vaginal microbiota prior to or during early pregnancy typically transition to a vaginal microbiota of *Lactobacillus*-dominance as gestation progresses (50, 51, 66), suggesting that pregnancy entails selective pressures for *Lactobacillus*-dominance in the vaginal microenvironment. Therefore, it is important to know if the vaginal microbiota of the mouse has similar or ecologically parallel characteristics to those of the human vagina microbiota given the propensity for the use of the mouse model in studies of pregnancy, intrauterine infection, and spontaneous preterm birth.

**Table 1.** Description of previous 16S rRNA gene studies of the pregnant mouse microbiome.

| Author | Year | Body site | Microbiota culture methods | Mouse strain | Key microbiota findings |
| --- | --- | --- | --- | --- | --- |
| Gohir, et al. (119) | 2015 | Intestine | Not performed | C57BL/6J | *Akkermansia*, *Clostridium*, *Bacteroides*, and *Bifidobacterium* were increased in pregnant control mice compared to non-pregnant mice. Other abundant taxa observed in pregnant mice included *Lactobacillus*, *Alistipes*, and *Lachnospiraceae*. |
| Jašarević, et al. (92) | 2017 | Intestine, vagina | Not performed | C57BL/6:129 | Relatively abundant taxa in the intestine included S24-7, *Prevotella*, an unclassified Clostridiales, and *Sutterella*. In the vagina, the top mean relative abundant taxa were *Aggregatibacter*, *Lachnospiraceae*, and Clostridiales at embryonic day 7.5. |
| Nuriel-Ohayon, et al. (120) | 2019 | Intestine | Not performed | Swiss Webster | Most abundant taxa included S24-7, Clostridiales, Rikenellaceae, Bifidobacteria, *Lachnospiraceae*, *Lactobacillus*, and *Turicibacter*. |
| Younge, et al. (93) | 2019 | Intestine, vagina | Not performed | C57BL/6 | Most abundant taxa in the stool included S24-7, *Candidatus* *Athromitus*, and *Allobaculum*, while the vagina was largely dominated by *Kurthia*. |
| Faas, et al. (39) | 2020 | Intestine | Not performed | C57BL/6J OlaHsd | The intestinal microbiota at gestational days 7 and 14 were similar to the microbiota before pregnancy, however at gestational day 18, the microbiota became less diverse and were predominantly dominant in *Allobaculum*. |
| Theis, et al. (21) | 2020 | Intestine, lung, oral cavity, vagina | Homogenized tissue or Eswab fluid was plated onto tryptic soy agar with 5% sheep blood and Chocolate agar plates under anoxic, hypoxic (5% CO<sub>2</sub>, 5% O<sub>2</sub> and 90% N<sub>2</sub>), and oxic conditions at 37° C for 7 days. | C57BL/6 | Several bacterial taxa abundant in the intestine included S24-7, *Candidatus* *Athromitus*, *Bacteroides*, *Helicobacter*, and *Lactobacillus*. The lung was predominantly S24-7 and *Lactobacillus*. The oral cavity was abundant in *Streptococcus*, *Lactobacillus*, and Pasteurellaceae, while the vagina was almost exclusively Pasteurellaceae and *Helicobacter*. |
| Liu, et al. (121) | 2021 | Intestine | Not performed | ICR | Abundant taxa in the control group included Prevotellaceae and *Acinetobacter* and were primarily composed of members of Firmicutes and Bacteroidetes phyla |

The microbiota of the oral cavity, lung, and intestine can also influence human pregnancy outcomes. Several studies have detected microorganisms from the oral cavity of pregnant women, especially *Fusobacterium nucleatum*, in the amniotic cavity, which is presumably due to hematogenous transfer and can result in stillbirth or spontaneous preterm birth (67, 68). *Mycoplasma pneumoniae* and *Mycobacterium tuberculosis*, bacteria known to colonize the human lung, have also been implicated in human intra-amniotic infection (69, 70). Additionally, *Streptococcus agalactiae* is a commensal bacterium in the human intestine and vagina, however, colonization of the neonate by this bacterium during delivery can cause adverse neonatal outcomes such as sepsis (71–73). Given the potential for pregnancy complications caused in part by microorganisms from the oral cavity, lung, intestine, and vagina in humans, understanding the structure of these murine microbiotas during gestation is required if the mouse is to be effectively used as a model for investigating the role of the microbiota in obstetrical complications.

The objectives of this study were therefore to characterize the microbiotas of the oral cavity, lung, intestine, and vagina of the pregnant mouse using anoxic, hypoxic and oxic culture, as well as 16S rRNA gene sequencing, and to compare and contrast the effectiveness of these different microbiological approaches for characterizing the mouse microbiota **(Figure 1)**. We found variation by atmosphere in the composition of microorganisms cultured, with a greater diversity of bacteria recovered under anoxic conditions. However, it was the profiles of bacterial communities cultured under hypoxic and oxic conditions that best matched the structure of the 16S rRNA gene profiles of sampled body sites. Each body site had a unique microbiota; yet, multiple taxa were shared across body sites, suggesting a degree of interconnectedness among the microbiotas at these sites. Notably, potentially analogous to the human vaginal microbiota, the microbiota of the pregnant mouse vagina clustered into community state types (CSTs) based primarily on the dominance of two congeners, *Rodentibacter pneumotropicus* and *R*. *heylii*. Whole genome sequencing of cultured isolates of these two *Rodentibacter* species revealed genes associated with the utilization of glycogen, the predominant carbohydrate in the vagina. Importantly, the profiles of bacterial communities cultured from the vagina overlapped tightly with the 16S rRNA gene profiles of this body site. Therefore, culture can be used to accurately characterize the microbiota of the pregnant mouse vagina, and such can be successfully cultured and maintained in the laboratory and tractably manipulated for experimental *in vivo* studies of the vaginal microbiota and its role in pregnancy complications.

**Figure 1.**
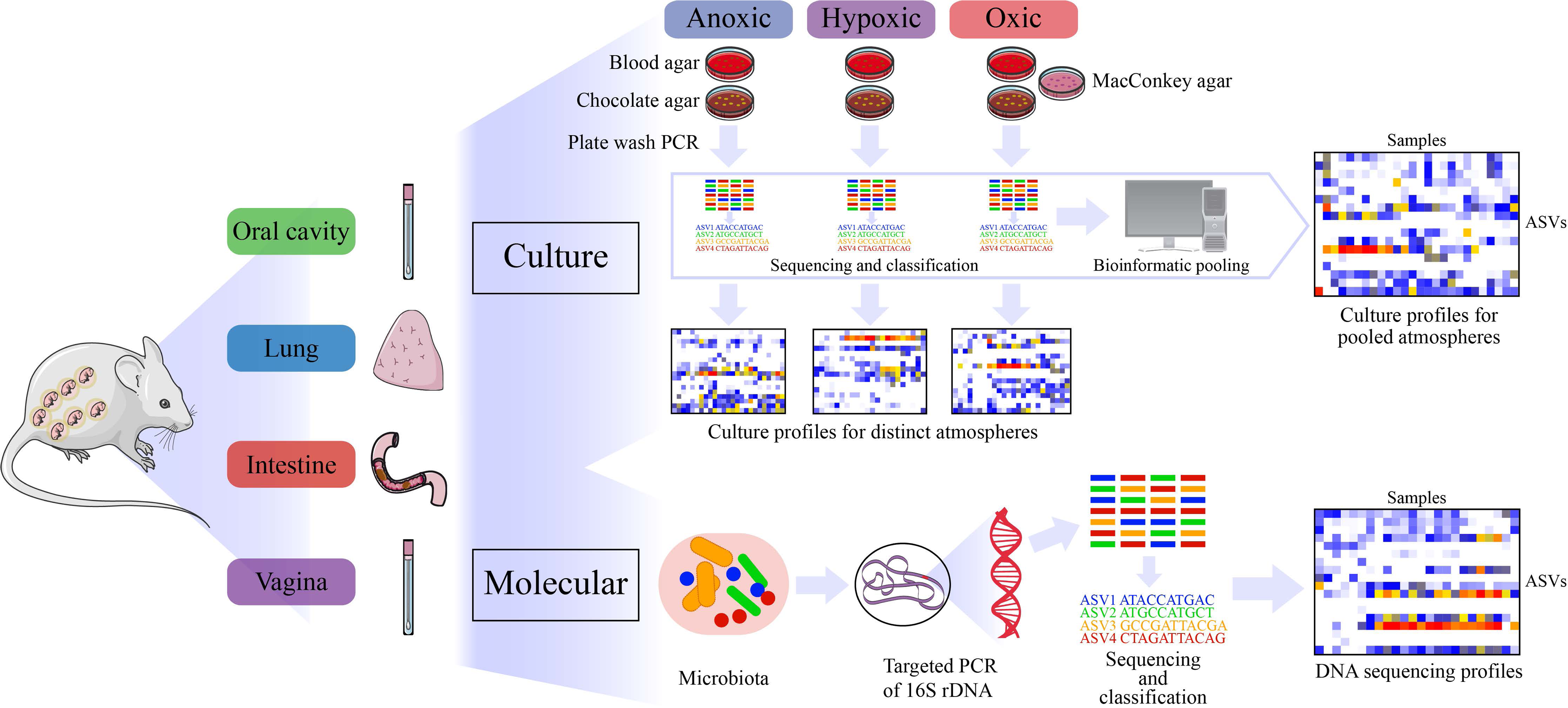
Study design for characterizing the microbiota of the oral cavity, intestine, lung, and vagina of pregnant mice. Briefly, two sets of samples were collected from each body site of 11 pregnant mice. One set of samples was used for culture and the other for molecular surveys. Cultures were performed on samples from each body site, under three different atmospheric conditions on multiple media types. Bacterial growth from each plate type was collected by plate washing with sterile PBS and then combined under each atmosphere. These samples subsequently had their DNA extracted followed by 16S rRNA gene amplification and sequencing. After classification of 16S rRNA gene sequences through DADA2, culture profiles for each body site under each atmosphere were generated as well as overall body site culture profiles after pooling the sequence data from all three atmospheres. Samples for molecular surveys had their DNA extracted directly from the samples followed by 16S rRNA gene amplification, sequencing, and classification to generate molecular profiles.

## RESULTS

### Influence of atmosphere and body site on the microbiota cultured from the oral cavity, lung, intestine, and vagina

#### Alpha diversity

Alpha diversity (i.e., the diversity within a single community) of the cultured microbiota varied by atmosphere (i.e., anoxic, hypoxic, oxic conditions) in all body sites except the vagina **(Figure 2A-D)**. In general, the cultured microbiota under anoxic conditions were more diverse than the cultured microbiota under hypoxic and oxic conditions; this observation was most pronounced for the cultured intestinal microbiota **(Figure 2C)**. After bioinformatically pooling the cultured microbiota data from all atmospheres for each individual mouse by body site, variation in microbiota alpha diversity was clear among the four body sites **(Figure 2E)**. The cultured intestinal microbiota was consistently the most diverse, while the cultured vaginal microbiota was consistently the least diverse **(Figure 2E)**.

**Figure 2.**
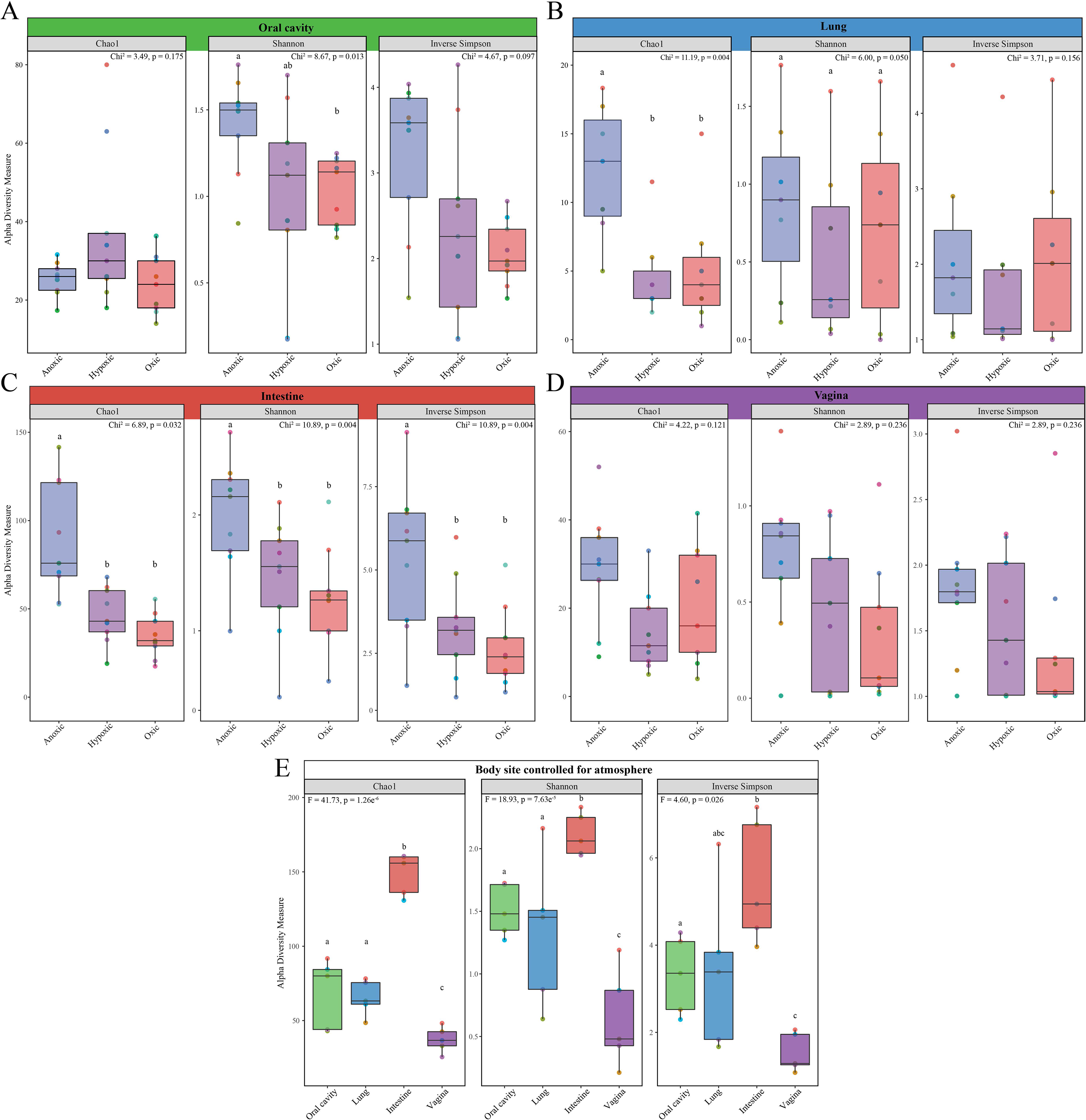
Alpha diversity comparisons between the microbiota cultured under different atmospheres for the oral cavity, lung, intestine, and vagina and between body sites. Bar plots indicate differences in three alpha diversity measures among anoxic, hypoxic, and oxic cultures of the oral cavity (A), lung (B), intestine (C), and vagina (D), as well as across body sites (E). For panel E, culture data from each atmosphere for each individual mouse by body site were bioinformatically pooled, and only mice with culture data from all body sites and all atmospheres (N = 5) were included in the analyses. Data points are color-coded by mouse ID and are consistent across panels. Lower case letters that are shared within each panel indicate pairwise comparisons that were not significant.

#### Beta diversity

Beta diversity (i.e., the diversity between two communities) of the cultured microbiota varied in composition and structure by both body site and atmosphere **(Table 2, Figure 3A-B, Supplemental Table 4-6)**. Although atmosphere was a global driver of variation of the cultured microbiota, when the data for each body site were assessed separately by atmosphere **(Supplemental Figures 1-4)**, variation in microbiota composition and structure was not observed for the lung or vagina **(Supplemental Table 5; Supplemental Figures 2 and 4)**. Notably, mouse identity contributed to variation of the bacteria cultured from the vagina but not to that of the bacteria cultured from the three other body sites. Additionally, vaginal samples appeared to cluster into six distinct groups based on the most abundant cultured taxa **(Supplemental Figure 4C)**, suggesting the existence of vaginal community state types in the mouse. After bioinformatically pooling the culture data from all atmospheres by body site for each mouse, mouse identity and body site were identified as primary drivers of variation in the cultured microbiota **(Figure 3)**, suggesting that the different atmospheres may have masked the influence of mouse identity in the previous analyses **(Table 3)**.

**Figure 3.**
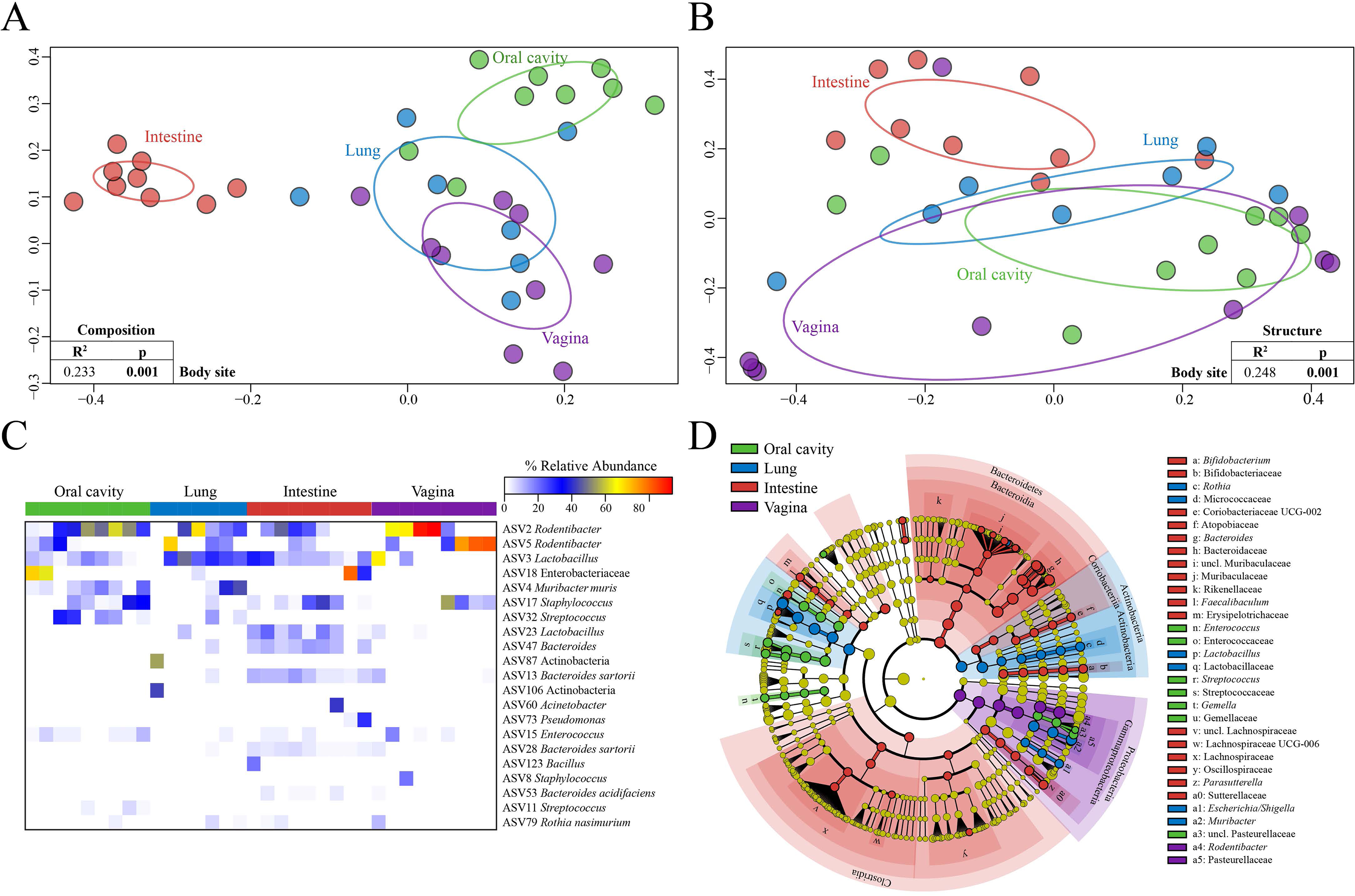
Comparisons of cultured microbiota from the oral cavity, lung, intestine, and vagina, controlled for atmosphere. Panels A and B contain Principal Coordinates Analysis (PCoA) plots illustrating variation among cultured microbiota of the oral cavity, lung, intestine and vagina using Jaccard dissimilarity index (A) for composition and Bray-Curtis dissimilarity index (B) for structure. Ellipses in (A) and (B) indicate standard deviation. The heatmap in panel C includes ASVs ≥ 1% average relative abundance within a single body site and samples are clustered by Bray-Curtis similarities within each body site. In panel D, LEfSe analysis identified taxa preferentially recovered in a particular body site.

**Table 2.**
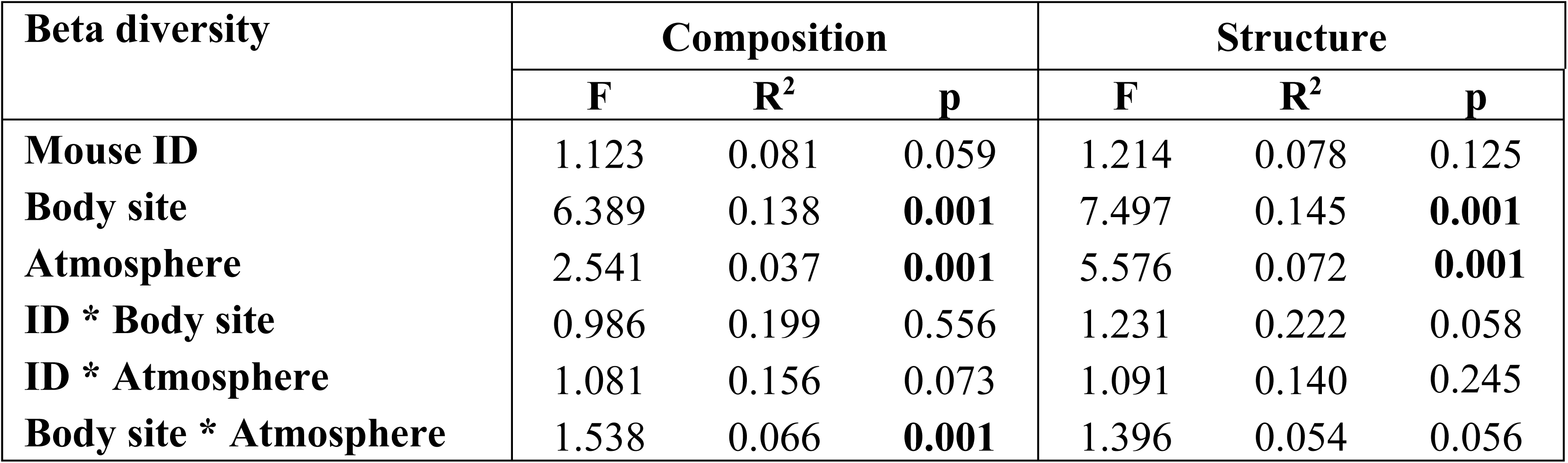
Global comparisons of the cultured murine microbiota.

**Table 3.**
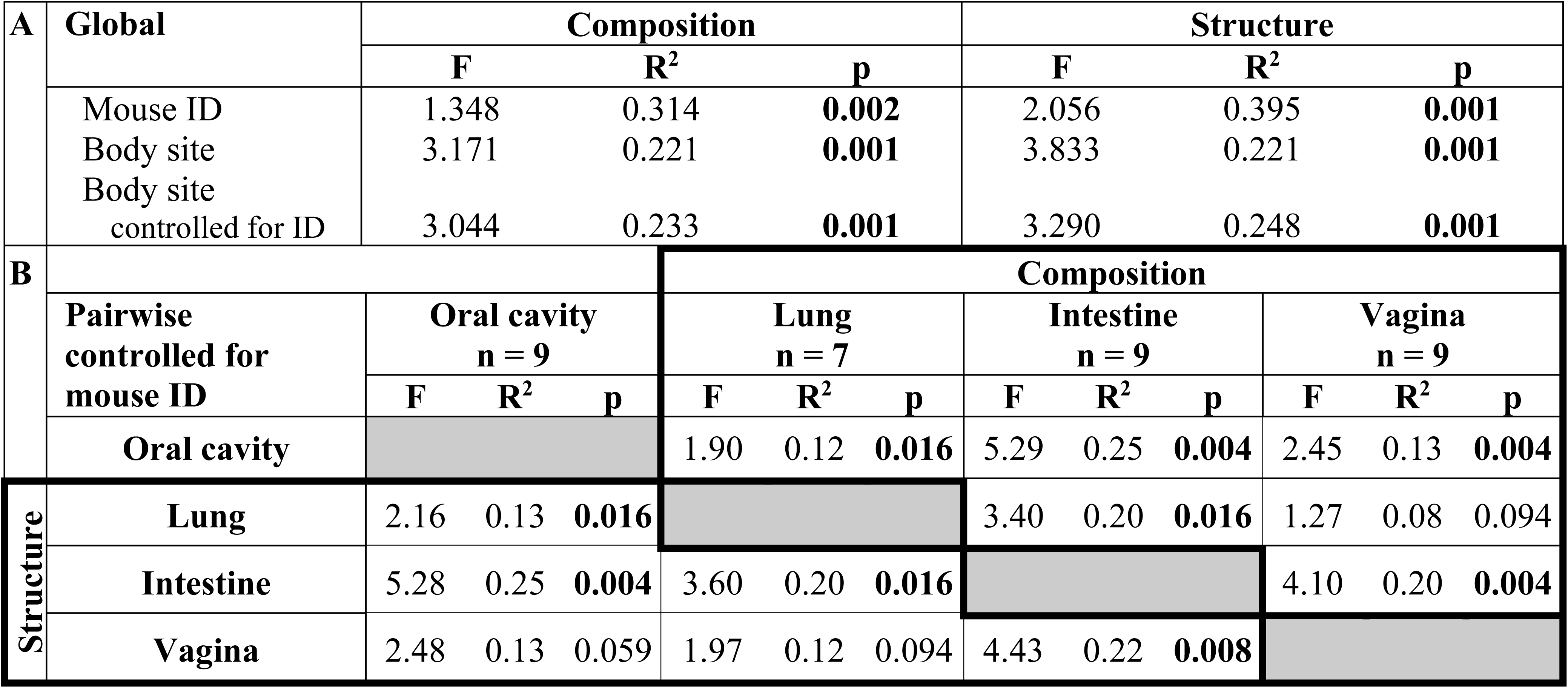
Global (A) and pairwise (B) comparisons of the cultured murine microbiota after bioinformatically pooling data across atmospheres by body site for each individual mouse.

### Influence of atmosphere, controlled for body site, on the cultured microbiota

#### Oral cavity microbiota preferentially recovered under different atmospheres

Under anoxic conditions, cultures of oral cavity microbiota appeared to cluster based on the relative abundance of either 1) *Lactobacillus* (Amplicon Sequence Variant 3 or ASV 3), *Muribacter muris* (ASV 4), and *Streptococcus* (ASV 32), or 2) *Rodentibacter* (ASV 2) and *Staphylococcus (*ASV 17) **(Supplemental Figure 1C)**. This was contrasted with the cultures recovered under hypoxic and oxic conditions, which were consistently dominated by *Muribacter muris, Rodentibacter*, and *Staphylococcus* **(Supplemental Figure 1C)**. Two LEfSe analyses were performed, one that was not restricted to a particular taxonomic classification level (i.e., hierarchical analysis) and one that was restricted to the level of ASV. Hierarchical LEfSe analysis revealed preferential recovery of bacteria from the phylum Firmicutes under anoxic conditions, specifically of the genera *Enterococcus, Lactobacillus,* and *Streptococcus* **(Supplemental Figure 1D)**, while members of the phyla Proteobacteria and Actinobacteria were preferentially recovered under oxic conditions, including the genera *Rodentibacter* and *Rothia.* Specific ASVs of each of these genera were identified in the ASV-level analysis **(Supplemental Figure 5A)** and included prominent ASVs 2, 3, 15, 79, from 4 of the genera identified in the hierarchical analysis **(Supplemental Figure 1D)**.

#### Intestinal microbiota preferentially recovered under different atmospheres

The microbiota cultured from the intestine under anoxic conditions were characterized by high relative abundances of several *Bacteroides* and *Lactobacillus* ASVs, as well as low relative abundances of *Bifidobacterium* and *Parasutterella* ASVs **(Supplemental Figure 3C)**. Hierarchical LEfSe analysis revealed a large number of taxa that were cultured preferentially under anoxic conditions compared to the other atmospheric conditions **(Supplemental Figure 3D)**. Notably, the phyla Bacteroidetes and Actinobacteria were heavily represented, as well as members of Firmicutes, especially Lachnospiraceae and Oscillospiraceae, and to a lesser extent members of the phyla “Desulfobacterota phyl. nov.” (74) (originally classified under the delta subdivision of Proteobacteria) and Verrucomicrobia. At the genus level, 13 genera were preferentially recovered in culture under anoxic conditions, including *Akkermansia, Bacteroides, Bifidobacterium, Colidextribacter,* Coriobacteriaceae UCG-002*, Desulfovibrio, Enterorhabdus, Faecalibaculum, Parasutterella*, *Lachnoclostridium,* Lachnospiraceae UCG-006, *Muribaculum,* and *Rikenella*. *Staphylococcus* was the only genus that was preferentially recovered under oxic conditions, and no genera were preferentially recovered under hypoxic conditions **(Supplemental Figure 3D)**. The trends in the hierarchical analysis were consistent with those in the analysis restricted to the ASV-level. With respect to the intestine, 18 ASVs were preferentially recovered under anoxic conditions, including *Akkermansia muciniphila,* multiple *Bacteroides* ASVs, *Bifidobacterium, Lactobacillus*, and *Parasutterella* **(Supplemental Figure 5C)**. *Bacteroides* and *Lactobacillus* were also recovered in culture under hypoxic and oxic atmospheric conditions, but *Bifidobacterium* and *Parasutterella* were not **(Supplemental Figure 3C)**. *Rodentibacter* and *Staphylococcus* ASVs constituted a large proportion of the cultures obtained under hypoxic and oxic conditions, yet they were not recovered under anoxic conditions **(Supplemental Figure 3C)**. The ASV-only analysis identified only one feature as discriminant of oxic and hypoxic cultures, a *Staphylococcus* (ASV 17) and *Bacteroides acidifaciens* (ASV 26), respectively **(Supplemental Figure 5C)**.

#### Lung and vaginal microbiota preferentially recovered under different atmospheres

The profiles of microbiota cultured from the lung and vagina were not affected by atmosphere **(Supplemental Figures 2, 4)**. However, LEfSe analysis identified *Streptococcus* (ASV 32) and *Bacteroides sartorii* (ASV 28) as being preferentially recovered under anoxic conditions from the lung **(Supplemental Figure 5B)**. No taxa or ASVs were identified as being differentially recovered based on atmospheric conditions from the vagina.

### Influence of body site, controlled for atmosphere, on the cultured microbiota

Between the four body sites, there were a total of 33 prominent ASVs (defined as having an average relative abundance ≥ 1% in at least one body site and atmosphere combination) **(Supplemental Figures 1C-4C)**. Five ASVs were prominent among all four body sites **(Supplemental Table 7)**. These five ASVs were classified as *Rodentibacter* (ASVs 2 and 5), *Lactobacillus* (ASV 3), *Staphylococcus* (ASV 17), and *Rothia nasimurium* (ASV 79). Twenty of the 33 ASVs were prominent in only one body site, typically either the intestine or lung, and limited to one or two samples at high relative abundance or multiple samples at a low relative abundance **(Supplemental Table 8)**.

*Rodentibacter* was cultured from nearly all vaginal samples and at high relative abundance, yet the presence and abundance of the two *Rodentibacter* ASVs differed among vaginal samples. Specifically, in most mice, only one of the two *Rodentibacter* ASVs were abundant **(Supplemental Figure 4C)**. In a minority of mice, both *Rodentibacter* ASVs were abundant. This contrasted with cultures from the other body sites, in which ASV 5 was much less common and ASV 2 was limited to recovery under hypoxic or oxic conditions, except in a few oral samples **(Supplemental Figures 1C-4C)**.

The prominent *Lactobacillus* (ASV 3) was cultured from most intestinal and lung samples regardless of atmosphere, exclusively under anoxic conditions from most of the oral samples and was highly abundant in only a single vaginal sample (the only vaginal sample without *Rodentibacter*). *Staphylococcus* (ASV 17) was commonly cultured from oral and intestinal samples but only rarely from lung samples. In the vagina, *Staphylococcus* (ASV 17) was exclusively cultured from samples that had an abundance of *Rodentibacter* ASV 5; it was only detected alongside *Rodentibacter* ASV 2 when ASV 5 was also abundant.

After bioinformatically pooling the culture data from each atmosphere by body site for each mouse, 21 ASVs were prominent in at least one body site **(Figure 3C)**. Three ASVs were prominent among all four body sites, with the *Rodentibacter* ASVs 2 and 5 having the greatest average relative abundance in vaginal cultures (36.1% and 33.2%, respectively), and ASV 3 (*Lactobacillus*) having the greatest average relative abundance in lung cultures (24.1%). Ten of these 21 ASVs were prominent in only one body site, with six being prominent only in intestinal cultures. Only one of the 21 prominent ASVs was exclusive to a single body site; ASV 106, an unclassified Actinobacteria, was unique to the lung.

LEfSe analysis revealed many taxa that were cultured preferentially from the intestine **(Figure 3D)**. Specifically, members of the phyla Bacteroidetes and Firmicutes were preferentially recovered in the intestine **(Figure 3D)**. At the genus level, 6 genera were preferentially recovered in cultures of the intestine, including *Bacteroides, Bifidobacterium,* Coriobacteriaceae UCG-002*, Faecalibaculum, Parasutterella*, and Lachnospiraceae UCG-006. Members of the Phylum Actinobacteria and genera *Escherichia/Shigella*, *Lactobacillus, Muribacter,* and *Rothia* were preferentially recovered from the lung. *Enterococcus, Streptococcus*, and *Gemella* were preferentially recovered from the oral cavity, while *Rodentibacter* was preferentially recovered from the vagina **(Figure 3D)**.

### Cultured microbiota contrasted with molecular characterizations of the same samples

Alpha diversity varied similarly for molecular profiles as was observed in the cultured microbiota. Variation was observed in both richness (Chao1, Friedman’s test: F = 16.91, p < 0.001) and evenness (Shannon and Inverse Simpson, Friedman’s test: F = 16.91, p < 0.001) between the oral cavity, intestine, and vagina. Pairwise comparisons revealed the intestine was more diverse than both the oral cavity (Wilcoxon singed rank tests for all three indices: W = 66, p < 0.001) and vagina (Wilcoxon singed rank tests for all three indices: W = 66, p < 0.001), while the oral cavity and vagina were not (Wilcoxon singed rank test: Chao1, W = 18, p = 0.206; Shannon, W = 40, p = 0.577; Inverse Simpson, W = 31, p = 0.898). The low alpha diversities were largely due to high relative abundances of *Streptococcus danieliae* (ASV1) and *Rodentibacter* (ASV 2 and 5) observed in the oral cavity and vagina, respectively **(Figure 4A)**.

**Figure 4.**
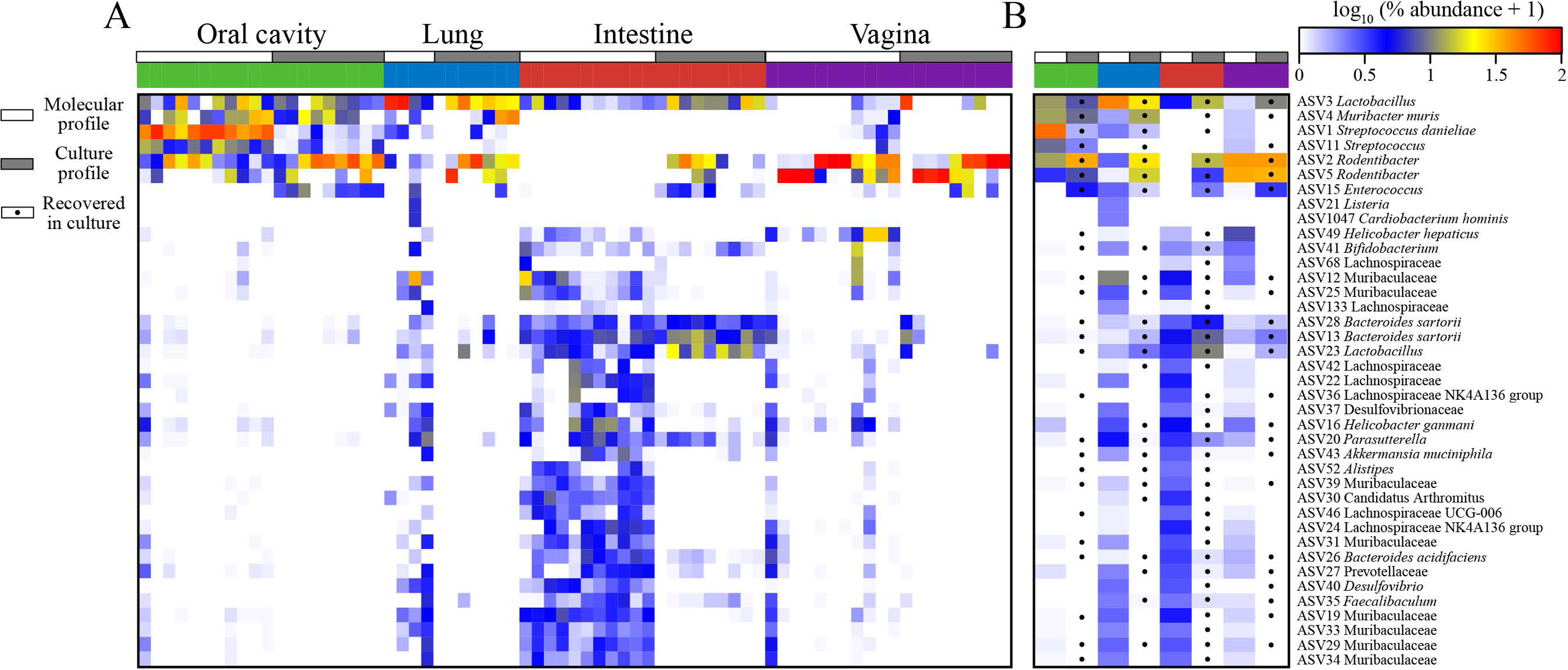
Comparisons of sequenced microbiota and cultured microbiota from the oral cavity, lung, intestine, and vagina. The heatmap in panel A represents log transformed precent relative abundances. In panel B, molecular and culture profiles were separately averaged with dots indicating whether an ASV was detected in culture. ASVs were included if they were ≥ 1% average relative abundance in the molecular profiles in one of the four body sites.

In total, cultured surveys accounted for 411 ASVs contrasted with 751 ASVs in molecular surveys **(Supplemental Table 9)**. Remarkably, only 339 ASVs were detected in both datasets, however both datasets had numerous ASVs not observed in the opposing dataset. For each body site, more ASVs were detected in molecular surveys than culture surveys except for the lung **(Supplemental Table 9)**. Of the prominent ASVs among both datasets (ASVs with an average relative abundance ≥ 1% in at least one body site from culture or molecular samples), most ASVs were detected in both datasets overall, over 90% (46/51), and at least half were observed in both culture and molecular datasets at each body site **(Figures 4A)**. Only 3 of the 39 prominent molecular ASVs were not detected in culture surveys **(Figure 4B)**. Two of these were only prominent in the lung, while the third, ASV 22 was prominent in the lung and intestine, and was detected in the intestine of all 11 mice. Of the 21 ASVs prominent in the cultured bacterial profiles **(Figure 4A)**, 11 were detected in all four body sites via molecular surveys while only two were not detected in any body site (ASV 106 and 123). Despite sharing a majority of prominent ASVs, correlations between culture and molecular profiles were only observed among the intestine and vagina **(Table 4)**, likely due to the overlap of prominent ASVs and the dominance of *Rodentibacter* ASVs in the vagina.

**Table 4.** Correlations between cultured microbiota recovered under anoxic, hypoxic, oxic atmospheres, or after pooling data from all three atmospheres, and molecular 16S rRNA gene profiles.

|  | Anoxic |  | Hypoxic |  | Oxic |  | Pooled Atmospheres |  |
| --- | --- | --- | --- | --- | --- | --- | --- | --- |
|  | *r* | *p* | *r* | *p* | *r* | *p* | *r* | *p* |
| Oral cavity | -0.1185 | 0.7717 | -0.5196 | 0.8655 | -0.123 | 0.7409 | -0.0662 | 0.6316 |
| Intestine | 0.4847 | 0.0616 | **0.3982** | **0.0486** | **0.6139** | **0.0123** | **0.5511** | **0.0212** |
| Vagina | **0.4564** | **0.0072** | **0.747** | **0.0003** | **0.7149** | **0.0003** | **0.6965** | **0.0007** |

Spearman rank correlation coefficients (*r*) and corresponding *p* values are provided. The lung could not be assessed due to low sample size.

### Comparative genomics of the two predominant vaginal bacteria

The distinct distribution and relative abundance patterns of ASV 2 and ASV 5 in the bacterial profiles of vaginal samples warranted further investigation of their genomic potential. ASV 2, identified as *Rodentibacter pneumotropicus* by 16S rRNA gene BLAST (75) analysis of sequenced isolates, and ASV 5, identified as *Rodentibacter heylii*, were submitted for whole genome sequencing to assess how the genomic and functional features of these two distinct *Rodentibacter* isolates might explain their distribution and abundance patterns in the murine vagina. The assembled genomes of both isolates were incorporated into a phylogenomic analysis of *Rodentibacter* type strains **(Figure 5A)**, as well as all available *Rodentibacter* spp. genomes **(Figure 5B)**. The genomes of the ASV 2 and ASV 5 isolates clustered as expected, based on the 16S rRNA gene analysis, with the genomes of their conspecifics, and a summary of the general genomic features of the isolates and two additional strains is provided in **Supplemental Table 10**.

**Figure 5.**
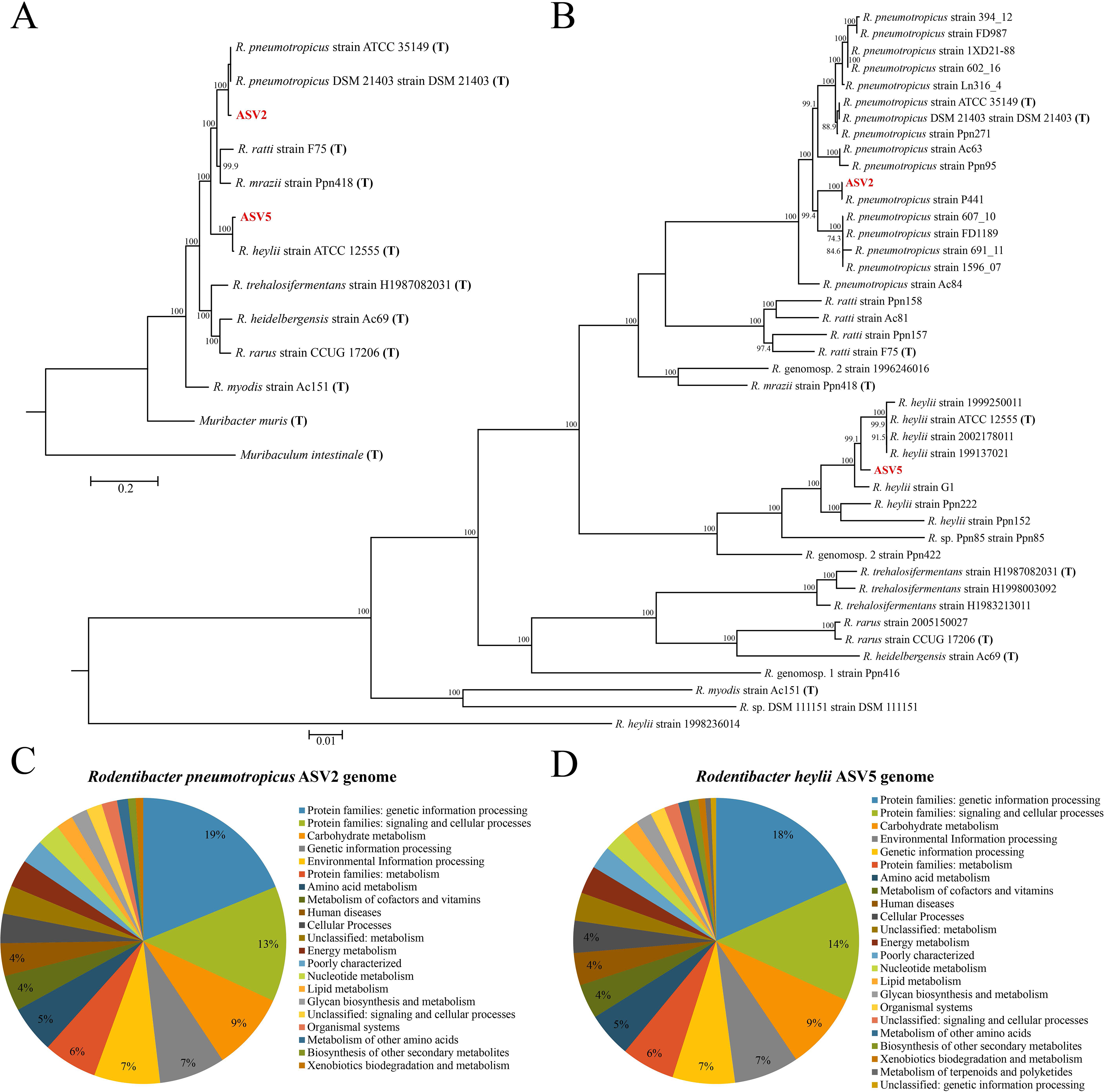
Phylogenomic and KEGG analysis of two vaginal *Rodentibacter* isolates. In panel A, the phylogenomic tree includes the *Rodentibacter* isolates ASV 2 and ASV 5 and all *Rodentibacter* type strains and in panel B, all published *Rodentibacter* genomes are included. Panels C and D summarize the distribution of functional KEGG pathways enriched in the genomes of the two isolates. Phylogenomic trees were constructed by comparing 92 conserved bacterial genes as described by Na et al. (112).

Of the 2,384 genes present in the ASV 2 genome, 1,505 could be confidently assigned to a KEGG molecular network (76). The most represented categories were genetic information processing, environmental information processing, and carbohydrate metabolism **(Figure 5C)**. Analysis of complete pathways for carbohydrate degradation indicated that ASV 2 has the capacity to utilize glycogen and 12 sugars: 2-deoxy-alpha-D-ribose-1-phosphate, D-arabinose, fructose, fucose, galactose, glucose, D-mannose, melibiose, ribose, trehalose, xylose and nine-carbon keto sugars (sialic acids N-Acetylneuraminate and N-acetylmannosamine). The genomic potential of ASV 2 was compared to that of 16 other reported *Rodentibacter pneumotropicus* strains for which published genomes were available. The published strains contained 1,565 core genes that were also present in ASV 2. Based on Prokka annotation of genomes (77), the pangenome of the 17 strains consisted of 4,389 genes, with each strain containing an average of 2,178 genes. Notably, ASV 2 contained the most genes (2, 321) among these strains, followed by *R. pneumotropicus* strain Ac84 (2, 311). Compared to the other *R. pneumotropicus* genomes, the genome of ASV 2 contained 83 unique genes, of which 81 are hypothetical proteins. The two unique genes with annotated functions were identified as DNA (cytosine-5-)-methyltransferase (*ydiO*) and serine/threonine-protein phosphatase 1 (*pphA*). An additional 25 annotated genes were unique to ASV 2 and its most phylogenetically similar strain P441, including a secretory immunoglobulin A-binding protein (*esiB*), bifunctional polymyxin resistance protein (*arnA*), and lipooligosaccharide biosynthesis protein lex-1 (*lex1*).

For the *R. heylii* isolate ASV 5, 1,537 of 2,474 genes were confidently assigned to a KEGG molecular network (76), and as with ASV 2, genetic information processing, environmental information processing, and carbohydrate metabolism were the most represented categories **(Figure 5)**. Complete pathways for carbohydrate degradation were very similar to those for ASV 2, including glycogen metabolism, with the exception that ASV 5 is not able to degrade 2-deoxy-alpha-D-ribose-1-phosphate, and it is able to degrade both L- and D- arabinose isomers, whereas ASV 2 can only utilize D-arabinose. The previously published genomes of 7 *R. heylii* strains have a core genome of 1,649 genes, of which 1,644 were present in the genome of ASV 5. The 5 missing genes included two hypothetical proteins, Lipopolysaccharide export system permease protein LptG (*lptG*), a duplicate outer membrane protein A (*ompA*), and a duplicate Anthranilate synthase component 2 (*trpG*). Compared to the other *R. heylii* genomes, ASV 5 contained 182 unique genes, of which 155 were hypothetical proteins. Notable genes unique to ASV 5 include mRNA interferase toxin RelE (*relE*), a duplicate Lysozyme RrrD (*rrrD*), very short patch repair protein (*vsr*), Enterobactin exporter EntS (*entS*), a duplicate Endoribonuclease ToxN (*toxN*) found in only one other strain, and Colicin V secretion protein CvaA (*cvaA*). A unique feature of the genome of ASV 5 compared to those of other published *R. heylii* strains is the presence of genes from the lsr operon, which regulates the autoinducer-2 quorum sensing pathway, suggesting that this strain may exhibit quorum-sensing, which may partially contribute to the distinct community structures observed in the present study.

Several differences in metabolic pathways were evident between the genomes of ASVs 2 and 5. As facultative anaerobes, the genomes of ASVs 2 and 5 encode genes for fermentation; however, only ASV 2 has the necessary alcohol dehydrogenase gene, *adhE*, for metabolizing ethanol. Other features unique to ASV 2 include metabolism of nucleotide monophosphates, the amino acids alanine and proline, and the reduction of glutathione. Notably, ASV 2 is missing several enzymes involved in the TCA cycle including citrate synthase; conversely, ASV 5 is not. However, this observation was not unique to ASV 2, as these enzymes are also missing from the other published *R. pneumotropicus* genomes. Collectively, they encode for and use citrate lyase as an alternative route for citrate degradation. Pathway features that are present in ASV 5 and yet missing in ASV 2 include lysine decarboxylase (needed for the biosynthesis of cadaverine), prepilin peptidase (involved in pilus formation), nitrite reductase (involved in denitrification), UDP-glucose:undecaprenyl-phosphate glucose-1-phosphate transferase (involved in colanic acid synthesis), and several enzymes necessary for the biosynthesis of the sialic acid CMP-N-acetylneuraminate. One interesting metabolic difference between the two ASVs is that ASV 5 contains three genes for the degradation of glycogen whereas ASV 2 only contains one. Also, ASV 5 contains a suite of tight adherence protein genes (*tadB* and *tadD-G*) and several, but not all, genes necessary for operation of the type IV secretion system; *virB2* and *virB7* were not identified in the genome. Lastly, although the genomes of both isolates contain the gene encoding the LuxS protein (a metabolic protein also utilized in quorum sensing), only isolate ASV 5 carries the necessary downstream genes for quorum-sensing, suggesting a substantial ecological distinction between the two isolates.

Shared features of the genomes of both isolates involved in interacting with the extracellular environment include genes for the sec-SRP and tat export pathways, lap adhesins, type VI secretion system, and the metabolism of urea. Also, although the genome of ASV 2 does not have a putative prepilin peptidase gene, both isolates contain multiple genes involved in pilus formation. While both isolates share several notable functions associated with interacting and persisting in the environment, ASV 5 has a greater capacity to interact with the environment. The more robust genome of ASV 5 and the differences in metabolism warrant further exploration, as do the number of hypothetical proteins observed in the genomes of both species. Detailed experimental studies may elucidate the mechanisms underlying the distinct colonization patterns we observe in the mouse vagina, especially in the context of ASV 5’s apparent unique quorum-sensing ability.

## DISCUSSION

### Principal findings of the current study

Preferential recovery of cultured microbiota was observed between anoxic, hypoxic, and oxic atmospheres with greater diversity of bacteria recovered under anaerobic conditions for each body site except for the vagina. Diversity of cultured microbiota varied by body site with the intestine having the greatest and the vagina having the lowest bacterial diversity. While some variation was evident between the cultured microbiota and molecular surveys for each body site, there was a strong positive correlation between the cultured microbiota and molecular profiles of the vagina. Bacterial profiles of the vagina were dominated by one or two distinct *Rodentibacter* strains (ASVs 2 and 5) using both culture and molecular approaches, indicating that the culture approaches employed herein accurately captured the vaginal microbiota. Whole genome sequencing of these *Rodentibacter* strains identified many shared genomic features, including the ability to metabolize glycogen, yet there were also strain-specific features, most notably a suite of quorum sensing genes exclusively observed in the ASV 5 strain.

### Impacts of atmosphere on the cultured microbiota of the mouse

Bacteria are capable of growth and reproduction in a variety of atmospheric conditions but are often broadly categorized by their ability or lack thereof to utilize O_2_ as a terminal electron acceptor during aerobic respiration under oxic conditions (78, 79). Notably, most of the body sites that are the focus of this study are typically low in O_2_ concentration compared to ambient atmospheres and are thus often considered anaerobic environments (7). However, these sites exhibit an O_2_ gradient, as O_2_ diffuses out from the host tissues into the mucus layer and the tissue-microbiota interface (79). It has therefore been suggested that microbial culture at low level O_2_ concentrations (i.e., hypoxic atmospheric conditions) will facilitate the growth of bacteria present at this interface which are able to grow but are typically outcompeted by other bacteria at lower (anoxic) or higher (oxic) oxygen concentrations (i.e., the atmospheric conditions most frequently used for microbial culture) (79).

In the current study, anaerobic culture yielded the greatest diversity of bacteria for the intestine, lung, and oral cavity, but not for the vagina. This may suggest a bias of culturing anaerobic bacteria from the intestine, lung, and oral cavity or merely a greater capacity for anaerobic bacteria from these sites to grow under laboratory conditions. Regardless, the low degree of correlation between the culture and molecular profiles of the microbiotas in the oral cavity indicate that the culture methods used in this study were not sufficient for capturing the breadth of bacteria present in this body site. Notably, however, the cultured and molecular surveyed microbiotas of the vagina were largely congruent, especially when culture was performed under hypoxic conditions. This leads to two important conclusions. First, when culturing the vaginal microbiota of the pregnant mouse, culture under hypoxic conditions alone appears sufficient for capturing its members – oxic and anoxic cultures would only need to be performed if specific hypotheses about the microbiota and vaginal oxygen levels were being investigated. Second, the current study demonstrates that the vaginal microbiota of the pregnant mouse can be reliably captured through laboratory culture and thus it is feasible and tractable to generate culture libraries that can be used for *in vitro* and *in vivo* manipulative experimentation of the vaginal microbiota and/or intra-amniotic infection in murine animal models of pregnancy complications.

### Prior reports of the oral cavity, lung, and vaginal microbiotas of non-pregnant mice

The microbiotas of body sites other than the intestine in laboratory mice have been only infrequently characterized by 16S rRNA gene sequencing. Studies characterizing the microbiotas of the oral cavity, lung, or vagina of normal non-pregnant mice are identified and summarized in Supplemental Tables 1-3.

Most studies characterizing the microbiota of the murine oral cavity have focused on a single mouse strain (i.e., C57BL/6) **(Supplemental Table 1)**. The genera within the oral microbiota often differed between studies, suggesting that environment plays a large role in the composition of the oral microbiota. This was demonstrated explicitly when the oral microbiota of mice from different laboratories were compared (80). Of the relatively abundant genera in the oral cavity, *Lactobacillus, Staphylococcus,* and *Streptococcus* were observed in multiple studies (80–83). Notably, no studies have characterized the oral microbiota of mice using culture.

The microbiota of the murine lung has been characterized through several studies comparing the microbiota of diseased or treatment groups to that of control mice, as opposed to strictly descriptive studies of control or healthy mice **(Supplemental Table 2)**. Little overlap of abundant genera has been observed among studies. In fact, one study acquired mice from two different breeding facilities, characterized the microbiota of the lung, and found that there were no core bacteria common to all mice and not a single bacterium was shared between the majority of mice (84). Yet, the authors did observe convergence of the lung microbiota of mice acquired from different facilities after a week of cohabitation, suggesting that the lung microbiota is dynamic and largely influenced by housing and social environments. Despite the pronounced role of the environment on the lung microbiota, several bacterial genera were relatively abundant in multiple studies: *Streptococcus*, *Lactobacillus, Pseudomonas,* and *Staphylococcus* (5, 84–87). Two studies of the lung microbiota have utilized culture alongside molecular approaches. In the first, only one bacterium was recovered, *Micrococcus luteus,* and only from culture (5). In the second, *Stenotrophomonas* and *Ochrobactrum* were detected in both culture and molecular surveys of the lung (88).

Studies characterizing the vaginal microbiota of mice have also varied in the abundant genera observed, however, like the microbiotas of both the murine oral cavity and lung, members of *Lactobacillus, Staphylococcus,* and *Streptococcus* were observed in multiple studies **(Supplemental Table 3)**. Two studies have each observed mice with similar vaginal microbiotas that could be clustered into at least two Community State Types (CSTs). In the first study, vaginal microbiota samples could be clustered into two CSTs based largely on the relative abundance of *Streptococcus* (>50% in one group and ≤ 10% in the other) (5). The second study included five vaginal CSTs which were defined by varying relative abundances of *Staphylococcus, Enterococcus, Lactobacillus*, and multiple lower abundance taxa (89). Although no study has characterized the vaginal microbiota in mice using both culture and molecular methods, two older studies did perform culture-based characterization of the vaginal microbiota in mice (90, 91). Both studies cultured members of *Streptococcus, Staphylococcus,* and *Lactobacillus*; one also consistently recovered *Corynebacterium* and *Actinomyces* (90), and the other recovered members of the Enterobacteriaceae and Bacteroidaceae families (91).

### Prior reports of the oral cavity, lung, intestinal, and vaginal microbiotas of pregnant mice

Excluding our current and prior study (21), the data of which overlap, the mouse intestinal microbiota during pregnancy has been characterized six times, and the vaginal microbiota has been characterized twice **(Table 1)**. Among the studies that characterized the intestinal microbiota of pregnant mice **(Table 1)**, approximately 18 bacterial taxa were observed at high relative abundances. The following taxa were observed at high relative abundances in multiple studies: S24-7, *Allobaculum*, *Bacteroides*, *Bifidobacterium*, Candidatus Athromitus, Clostridiales, *Lactobacillus*, and Lachnospiraceae.

The two prior studies which characterized the vaginal microbiota of pregnant mice also simultaneously characterized that of the intestine (92, 93). In the first study, researchers investigated the effect of stress on these microbiotas and subsequent downstream effects on the microbial colonization of newborn mice. 16S rRNA gene sequencing was performed on maternal fecal samples collected daily and on vaginal fluid collected on embryonic day 7.5 (92). The bacterial taxa that were relatively abundant in the fecal samples included *Sutterella, Prevotella*, S24-7, *Bacteroides, Odoribacter*, Desulfovibrionaceae, Lachnospiraceae, Ruminococcaceae, and *Oscillospira*. The alpha diversity of the maternal intestinal microbiota decreased early in pregnancy and the composition of the microbiota differed between early and late pregnancy. The vaginal microbiota of the pregnant control mice at 7.5 days gestation were mainly composed of Clostridiales, *Aggregatibacter*, Lachnospiraceae, *Prevotella, Helicobacter,* and S24-7.

In the second study, researchers evaluated the fetal compartments of mice for evidence of *in utero* bacterial colonization and characterized maternal intestinal and vaginal microbiotas to assess the source of any potential bacterial signals detected in the fetus (93). Samples from the maternal stool were relatively abundant in Candidatus Arthromitus, S24-7, and *Lactobacillus*, while the vaginal samples were predominantly composed of *Kurthia gibsonii*.

In the current study, similar to prior studies **(Table 1)**, we observed high relative abundances of Muribaculaceae (i.e., S24-7), *Bacteroides*, *Bifidobacterium*, Desulfovibrionaceae, *Lactobacillus*, and Lachnospiraceae in the intestinal microbiota of the pregnant mouse **(Figure 4)**. Yet, our findings for the vaginal microbiota were distinct. We found the vaginal microbiota of pregnant mice to be dominated by *Rodentibacter*, *Helicobacter*, and *Lactobacillus* **(Figure 4)**. The differences between the microbiota observed by Jašarević et al. (92) and those described in current study could be due to the gestational age at time of sampling. In the former, samples of the vaginal microbiota were taken during early gestation, embryonic day 7.5 (E7.5) whereas in the present study they were taken at late gestation (E17.5), which may suggest a shift in the vaginal microbiota that occurs between early and late gestation. In Younge et al. (93), vaginal samples were collected at one of three timepoints (E14-16, E17-18, E19-20) between mid to late gestation, however only two mice were sampled per group. The vaginal microbiota of the earlier timepoint consisted primarily of Candidatus Arthromitus and S24-7, which was similar to what was observed in the stool samples from those same mice. At the latter two timepoints, the vaginal microbiota were distinct from maternal stool with low diversity and high relative abundance of *Kurthia gibsonii*. Although no *Kurthia* sequences were detected in the current study, the low diversity observed in both studies suggests that the murine vaginal microenvironment changes during pregnancy and is permissive to the dominance of certain bacteria in the vagina.

### Bacterial Community State Types (CSTs) of the mouse vagina

In a previous study, five Community State Types (CSTs) were suggested for the non-pregnant mouse vagina (89). The authors described these as being dominated by *Staphylococcus* and/or *Enterococcus*, *Lactobacillus*, or a mixed population of bacteria. While these bacteria were also detected in our study of pregnant mice, aside from *Lactobacillus*, they were not observed at high relative abundances, suggesting a potential shift in the vaginal microbiota of non-pregnant mice upon becoming pregnant. In our study, we almost exclusively observed a vaginal microbiota dominated by one or two distinct *Rodentibacter* strains that were widespread among the mice. These dominant strains/ASVs potentially mirror the dominance of *Lactobacillus crispatus* and *L*. *iners* in prominent CSTs of the human vagina (62), suggesting the vagina of the pregnant mouse may represent a similar but unique ecological niche conducive to the proliferation of only a few dominant bacteria. This is especially interesting considering that during human pregnancy the vaginal microbiota typically shifts even more dramatically to a *Lactobacillus-*dominant community, especially later in gestation, and for those women who had non-*Lactobacillus*- dominant communities before pregnancy (66).

### Novel insights into *Rodentibacter* strains in the pregnant mouse vagina

The assembled genomes for cultured isolates of *Rodentibacter* ASVs 2 and 5 are representative of *R. pneumotropicus* and *R. heylii,* respectively **(Figure 5)**. It is unclear if these strains are uniquely adapted to murine vaginal microenvironments in general, or if this phenomenon is limited to pregnant mice, or even potentially pregnant mice in the specific animal housing facility under investigation here. Notably, this may be a general phenomenon as Jasarevic et al. (94) recently found *Pasteurella pneumotropica* to be abundant in the vagina of mice after pregnancy. In 2017, *P. pneumotropica* was reclassified to the genus *Rodentibacter* (94), indicating that this observation of *Rodentibacter* dominance is not exclusive to our animal facility.

The functional potential of both species suggests a wide range of metabolic capabilities like the ability to process various sugar sources, which may partially explain why both isolates were detected in multiple body sites of the mouse. Additionally, the genomes of both isolates indicate these strains can utilize glycogen which is a primary carbon source in the vagina (95). The larger genome size of ASV 5 and the greater number genes encoding of glycogen degradation enzymes may provide a more robust capacity to colonize and persist in the vaginal microenvironment and may partially explain why other ASVs co-occurred less frequently in ASV 5-dominant vaginal samples than in ASV 2-dominant samples.

The relationship of these two *Rodentibacter* isolates and their murine host appears highly similar to the relationship between *Lactobacillus crispatus* and *L*. *iners* and their human host. First, members of both genera are capable of inhabiting multiple body sites of their hosts (96–101). Second, the highest relative abundance among the populated body sites in both hosts is within the vagina, wherein relative abundances can exceed 90% of the sequenced microbiota (50, 59, 62, 66). Third, both isolates have the genomic capacity to degrade glycogen, which is a predominant carbon source in the mammalian vagina and is associated with the abundance of *Lactobacillus* in the human vagina (60, 102). Although *Rodentibacter* dominance in the pregnant mouse vagina has not been previously documented, this microbiota has been understudied **(Table 1)**. It is possible that a low diversity microbiota dominated by *Rodentibacter* is evidence of a shift in vaginal microbiota structure during pregnancy in the mouse. In humans, *Lactobacillus*-dominance during normal pregnancy is common and associated with healthy term gestations, whereas more diverse vaginal microbiotas are less common among pregnant women and have been associated with adverse pregnancy outcomes (60, 66). The *Rodentibacter*- dominant vaginal microbiotas observed in this study may represent a similar transition in mice. Specifically, the vaginal microbiotas of mice may be typically more diverse, akin to CST IV in the human vagina, and transition to a less diverse, *Rodentibacter*-dominant state during pregnancy. This needs to be investigated further.

### Strengths of this study

This was the first study to simultaneously characterize the microbiota of the oral cavity, lung, intestine, and vagina of the pregnant mouse through both culture and 16S rRNA gene sequence-based approaches. It was also the first study to consider the extent to which culturing the microbiota from these body sites under different atmospheric conditions captured the complete site-specific microbiota, as defined through the molecular surveys. This study revealed strong associations of *Rodentibacter* strains with the vagina of the pregnant mouse, and whole genome sequencing of cultured representatives of these strains identified functional features that may explain their dominance with the murine vagina during pregnancy.

### Limitations of this study

This study focused on C57BL/6 late-gestation pregnant mice from a single facility. This is important because there can be variation in the microbiota of mice across facilities (103, 104), as well as in the same laboratories over time (19). Therefore, it is not yet clear the extent to which the patterns in body-site specific microbiota data reported herein can be extrapolated to other studies. Additionally, non-pregnant mice and samples from pregnant mice at different gestational ages were not included and therefore the relationship of the microbiota throughout gestation could not be assessed. Nevertheless, this study provides a detailed foundational knowledge, based on multi-atmospheric culture and DNA-based sequencing approaches, of the microbiotas of the oral cavity, lung, intestine, and vagina of the pregnant mouse, thereby setting the stage for additional investigations into the reproductive microbial ecology of the mouse.

### Conclusions

The microbiota of the pregnant mouse includes bacteria shared among the oral cavity, lung, intestine, and vagina. Yet, variation was evident in the microbiotas across body sites. Comparisons of culture and molecular microbiota profiles indicate that culture, especially hypoxic culture, effectively captured the microbiota of the vagina, but not necessarily of the other body sites. As the vaginal microbiota can be effectively cultured, moving forward it can be tractably used for *in vitro* and *in vivo* experimentation evaluating relationships between the vaginal microbiota and adverse pregnancy outcomes in mice. The vaginal microbiota of the pregnant mouse appears to be dominated by one or two *Rodentibacter* strains, similar to the two *Lactobacillus*-dominant CSTs (i.e., I and III) in the human vagina during pregnancy. Whole genome sequencing of the *Rodentibacter* strains dominating the pregnant mouse vaginal microbiota here revealed the capacity to metabolize glycogen, a principal carbon source in the mammalian vagina. This capacity is also possessed by human vaginal lactobacilli. These findings suggest the existence of ecological parallels between the vaginal microbiotas of mice and humans during pregnancy. These parallels and their relevance to host reproduction warrant further investigation.

## MATERIALS AND METHODS

### Study subjects and sample collection

Culture and DNA sequencing surveys of samples from the oral cavity, lung, intestine, and vagina of 11 pregnant mice that were included in our previous study evaluating the *in utero* colonization hypothesis (21) were here analyzed in-depth in effort to characterize and compare the composition and structure of the pregnant mouse microbiota across body sites **(Figure 1)**. This study includes previously unpublished information on the culture of microorganisms from these body sites across atmospheric and growth media conditions, as well as functional genomic information on the principal *Rodentibacter* species inhabiting the murine vagina. Animal procedures were approved by the Institutional Animal Care and Use Committee at Wayne State University (protocol 18-03-0584).

### Bacterial culture

Bacterial culture was performed on intestinal and lung tissues and oral and vaginal swabs under oxic, hypoxic (5% O_2_, 5% CO_2_), and anoxic (5% CO_2_, 10% H, 85% N) conditions at 37⁰C for seven days. Under each atmosphere, samples were plated in duplicate onto tryptic soy agar with 5% sheep’s blood and chocolate agar. Samples were also plated on MacConkey agar under oxic conditions. If bacterial growth was observed (most typically a lawn of bacteria or too many colonies to count), the bacteria were collected by pipetting 1-2 ml of sterile PBS solution onto the agar plate and dislodging colonies with sterile and disposable spreaders and loops. These plate wash solutions (105) were stored at −80⁰C until DNA extractions were performed. DNA extractions were completed using a Qiagen DNeasy PowerSoil (Germantown, MD) extraction kit, as previously described (21). The V4 region of the 16S rRNA gene copies in DNA extractions were targeted using protocols previously described in Kozich et al. (106), and sequenced on an Illumina MiSeq system at Wayne State University, as previously described in Theis et al. (21). Ultimately, 16S rRNA gene sequence libraries were generated for the cultures from 117/ 132 (89%) murine body site samples.

### DNA sequencing surveys

Tissue samples of the lung, distal intestine, and proximal intestine were collected in addition to swabs of the oral cavity and vagina and stored at −80⁰C until DNA extractions were performed. DNA extractions of samples for molecular surveys were performed in a biological safety cabinet by study personnel donning sterile surgical gowns, masks, full hoods, and powder-free exam gloves. Extracted tissue masses ranged from to 0.016 to 0.107 g, 0.053 to 0.097 g, and 0.034 to 0.138 g for the lung, distal intestine, and proximal intestine, respectively. Two types of negative technical controls were included in the DNA extraction and sequencing processes to address potential background DNA contamination: 1) sterile swabs as a negative control for body sites sampled with a swab (i.e., oral and vaginal sites), and 2) extraction tubes with no biological input as a negative control for body sites from which tissue was collected (i.e., proximal intestine, distal intestine, and lung).

DNA was extracted from tissues, swabs, and technical controls (i.e., swabs [n = 11] and blank DNA extraction kits [n = 23]) using the Qiagen DNeasy PowerLyzer PowerSoil kit with minor modifications to the manufacturer’s protocol. Specifically, samples were added to the supplied bead tube along with 400 µl of bead solution, 200 µl of phenol-chloroform-isoamyl alcohol (pH 7 to 8), and 60 µl of solution C1. Mechanical lysis of cells was done using a bead beater for 30 seconds. Following centrifugation, the supernatants were transferred to new tubes and 100 µl of solutions C2 and C3 in addition to 1 µl of RNase A enzyme were added and tubes were incubated for 5 minutes at 4⁰C. After centrifugation, supernatants were transferred to new tubes containing 650 µl of solution C4 and 650 µl of 100% ethanol prior to adding to the filter column, and 60 µl of solution C6 for elution. The lysates were loaded onto filter columns until all sample lysates were spun through the filter columns. Five hundred microliters of solution C5 was added to the filter columns and centrifuged for 1 minute, the flowthrough was discarded, and the tube was centrifuged for an additional 3 min as a dry spin. Finally, 100 µl of solution C6 was placed on the filter column and incubated for 5 min before centrifuging for 30s to elute the extracted DNA. Purified DNA was stored at −20⁰C until 16S rRNA gene sequencing. Amplification and sequencing of the V4 region of the 16S rRNA gene were performed at the University of Michigan’s Center for Microbial Systems as previously described (21), with library builds performed in triplicate and pooled for each individual sample prior to the equimolar pooling of all sample libraries for multiplex sequencing.

### 16S rRNA gene sequence processing of bacterial culture and molecular samples

Raw sequence reads were processed using the DADA2 package in R following the tutorial pipeline as described by Callahan et al. (107) with minor modifications. Specifically, length of the reverse read truncation length was increased from 160 to 200 (“truncLen=c(240, 200)”), the maximum expected errors in reverse reads was increased from 2 to 7 (“maxEE=c(2, 7)”), and for sample inference samples were pooled to increase sensitivity (“pool=TRUE”). Sequences were ultimately classified into amplicon sequence variants (ASVs) and taxonomically identified using the Silva rRNA database v 138.1 (108, 109). After processing 16S rRNA gene sequences through DADA2, any ASVs identified as mitochondria, chloroplasts, and those not assigned to a bacterial phylum were removed.

Following DADA2 processing and removal of non-bacterial 16S rRNA gene sequences, only samples with libraries of at least 100 quality-filtered sequences were analyzed. From the culture samples two samples fell below this threshold and were removed from subsequent analyses (1 anoxic mid-intestine sample and 1 hypoxic lung sample). For the molecular samples, all vaginal, oral cavity, proximal intestine, and distal intestine sequence libraries met this criterion, but only five lung libraries remained. The full dataset included 176 biological samples, 15 blank extraction kit controls, and 17 negative swab controls representing a total of 1138 ASVs.

To validate that the bacterial signals detected in the molecular surveys of mouse samples were legitimate, the composition and structure of the bacterial profiles of tissues and swabs were contrasted with those of blank (n = 15) and blank swab (n = 17) technical controls using the adonis function in the *vegan* package. For each body site, the composition and structure of bacterial profiles were distinct from those of applicable negative controls **(Supplementary Table 11)**.

#### Removal of background DNA contaminant ASVs through decontam

After establishing that the 16S rRNA gene profiles of the tissue and swab samples from the pregnant mice were distinct from those of negative controls, the tissue and swab datasets were separately analyzed with *decontam* (110) to identity ASVs that were likely background DNA contaminants. Histogram plots of the distribution of prevalence scores indicated a threshold of 0.8 would be appropriate for both datasets, thereby retaining a large percentage of ASVs (82% in the tissue dataset and 72% in the swab dataset). Between the two datasets, 209 ASVs were below the 0.8 threshold and identified as contaminants. 24 ASVs were not detected in any biological samples from the molecular surveys and 179 ASVs had an average relative abundance below 1% for each of the biological sample types. Three of the remaining 6 ASVs, *Ralstonia* (ASV 76)*, Streptococcus* (ASV 520), and a *Bacillus* (ASV 6), were detected as contaminants in both tissue and swab datasets. *Streptococcus* (ASV 11)*, Muribacter* (ASV 4), and *Rodentibacter* (ASV 5), the 3 remaining ASVs, had average relative abundances above 1% in at least one body site from the opposing sample type (e.g., ASV 5 was above 1% average relative abundance in vaginal swabs but identified as a contaminant by *decontam* from the tissue dataset) suggesting that they may be legitimate sequences and were not considered contaminants and were retained in subsequent analyses. To allow for comparisons of the molecular datasets with the culture datasets, the *decontam* results were contrasted with the culture data to ensure that ASVs abundant in culture surveys were not removed as contaminants due to the fact that they were recovered via culture (i.e., they were legitimate as they were cultured by us). ASVs classified as contaminants through *decontam* were retained in subsequent analyses if they either were above 1% average relative abundance in at least one cultured body site or cultured from at least 5 mice for a given body site. Thirteen additional ASVs met these criteria and ultimately 16 out of the 209 ASVs identified as potential contaminants by *decontam* were kept in the datasets for subsequent analyses.

To aid comparisons of culture and molecular datasets, the molecular 16S rRNA gene profiles of the proximal and distal portions of the intestine were assessed for differences in their bacterial profiles using the adonis function in *vegan*. No differences were observed for either composition (mouse identity, p = 0.19; intestine locale, p = 0.47) or structure (mouse identity, p = 0.14; intestine locale, p = 0.26), and these samples were bioinformatically pooled by mouse identity and considered “intestinal” samples from molecular surveys. These data were contrasted with those of cultures from the mid-intestine.

### Whole genome sequencing and genomic analysis of isolates ASV 2 and ASV 5

#### DNA extraction from bacterial isolates

The 16S rRNA gene sequences associated with ASV 2 and ASV 5 were queried against a BLAST database of 16S rRNA gene sequences from isolates recovered and preserved during the previous study (21). Isolates with a 100% match were recovered from frozen stocks by plating 80 µl onto the media and atmosphere they were originally recovered on (both isolates were recovered on chocolate agar plates and under hypoxic atmospheric conditions (5% O_2,_ 5% CO_2_, 90% N_2_) and incubated for 48 hours. Colonies were then collected using sterile inoculating loops into 500 µl of sterile PBS and centrifuged at 15,000 x g for 10 minutes. DNA extractions were performed using a DNeasy PowerLyzer PowerSoil Kit (QIAGEN, Valencia, CA, USA) and the first step included resuspension of the pelleted colonies in 500 µl of bead solution and adding the total resuspended volume to the bead tubes. Two additional modifications to the manufacturer’s protocol were made: (1) 100 µl of solutions C2 and C3 were combined into a single step followed by 5-minute incubation at 4⁰C and subsequent centrifugation; (2) In the final elution step, DNA was eluted with 60 µl of solution C6 rather than 100 µl to increase DNA concentration. Extracted DNA was then stored at 4⁰C until submission (< 48 hours) for whole genome sequencing (WGS).

#### Construction and sequencing of sample DNA libraries

Libraries were built using the Illumina DNA Prep protocol and Nextera DNA CD indexes (Illumina). Libraries were sequenced at the Perinatology Research Branch using iSeq 100 reagents (Illumina) with the iSeq 100 system (Illumina) and an output of 2 x 150 bp paired-end reads.

#### DNA sequence processing and genome assembly

Adapters from the raw sequence reads were removed using Trimmomatic (v0.35). Genome assembly was performed using SPAdes (v 3.12.0) through the web-based Galaxy platform (111) with default parameters, except that k-mer sizes of 27, 37, 47, 57, 67, 77, and 87 were used. Contigs of less than 200 bp were removed after assembly and the average coverage per contig was 429 for ASV 2 and 256 for ASV 5.

#### Phylogenomic analysis of ASV 2 and ASV 5 isolates in the context of other Rodentibacter genomes

Phylogenomic analysis was performed using the up-to-date bacterial core gene (UBCG) tool (112) for phylogenomic tree inference, which utilized 92 core genes to assess phylogenomic relationships. The assembled genomes for isolates of ASV 2 and ASV 5, along with published *Rodentibacter* type strain genomes (downloaded through NCBI GenBank’s ftp server), and secondarily with all published *Rodentibacter* genomes, were processed through the UBCG pipeline using default parameters in Virtualbox. For the phylogenomic tree featuring only published type strains, *Muribaculum intestinale* was included as an outgroup and *Muribacter muris* was included as a within-family (Pasteurellaceae) outgroup. Trees were rooted on the midpoint using FigTree (v1.4.4) (113).

#### Genome annotation and analysis of ASV 2 and ASV 5 isolates

Prior to genome annotation, the contigs for each assembled genome were reordered and aligned to their closest strain, *R. pneumotropicus* strain P441 and *R. heylii* strain G1 for isolates ASV 2 and ASV 5, respectively, using Mauve multiple genome alignment software (v 20150226) (114). The 16 published genomes of *R. pneumotropicus* along with isolate ASV 2 were annotated with Prokka (v 1.14.5) (77) and processed through Roary (v 3.13.0) (115) to establish a core genome for *R. pneumotropicus* and to subsequently assess the representation of this core genome in the genome of isolate ASV 2. This process was repeated for isolate ASV 5 and the 7 published genomes of *R. heylii*. Prokka and Roary tools were run through Galaxy with default parameters except that paralog genes were not split when using Roary. For functional and pathway analysis, the genomes of isolates ASV 2 and ASV 5 were annotated with NCBI’s Prokaryotic Genome Annotation Pipeline (PGAP) (116) tool using default parameters. These annotated genomes were submitted to the KEGG Automatic Annotation Server (KAAS) (117) for KEGG pathway enrichment analysis using only bacteria included in the representative set of “Prokaryotes” as the template data set for KO assignment and a bi-directional best hit assignment method. For detailed metabolic comparisons of complete pathways, Pathway/Genome databases (PGDBs) were generated using the assembled genomes for isolates ASV 2 and ASV 5 with Pathway Tools software (v 25.0) (118). Default parameters were used; however, several manual refinement steps were required per the User’s Guide, including assigning probable enzymes, modified proteins, predicting transcription units, and inference of probable transporters.

### Statistical analysis

#### Alpha diversity

The alpha diversities of bacterial culture profiles for each body site under each atmosphere were characterized by the Chao1 (richness), Shannon (evenness), and Inverse Simpson (evenness) indices and variation in diversity was assessed for culture profiles under each atmosphere separately for each body site. Then after bioinformatically pooling mice that had culture profiles from three atmospheres for all four body sites (n = 5), diversity was compared between body sites. When comparing all three atmosphere culture profiles, datasets for each body site were rarefied to their lowest read depths (oral cavity: 3976 reads, lung: 358 reads, intestine: 4745 reads, vagina: 7719 reads). When comparing diversity between body sites, each individual mouse’s pooled samples were rarefied to the lowest read depth (20,954 reads). Alpha diversities of sequenced microbiota (molecular profiles) of the oral cavity, intestine, and vagina were also compared in the same way and rarefied to the lowest read depth (810 reads); the lung was excluded from this analysis due to low sample size (n = 4). Rarefaction was performed in R with the *phyloseq* package. Alpha diversities were calculated and visualized in R using the *phyloseq* package and labeled in Adobe Illustrator. Alpha diversities were statistically evaluated with the *rstatix* package in R by repeated measures ANOVA or Friedman’s ANOVA followed by paired t-tests or Wilcoxon signed rank tests when appropriate.

#### Beta diversity

Statistical comparisons of beta diversities were performed using the *vegan* and *pairwiseAdonis2* packages in RStudio (v 1.3.1093) and R (v. 4.0.3). Non-parametric multivariate analysis of variance (NPMANOVA) tests were used to evaluate the composition and structure of bacterial profiles using the Jaccard and Bray-Curtis Dissimilarity Indices, respectively. For comparisons where variation by mouse identity was observed, it was secondarily controlled for using the “strata” term on mouse identity in adonis and pairwise.adonis2 functions in *vegan* and *pairwiseAdonis2* packages, respectively. The composition and structure of bacterial profiles were visualized with Principal Coordinates Analysis (PCoA) plots generated using the RAM package in R (v. 4.0.3).

#### Linear discriminant analysis Effect Size (LEfSe)

LEfSe analyses were performed to identify features, taxa (assessed as hierarchical analyses), or ASVs (assessed as ASV-only analyses) that were preferentially recovered in different atmospheres for each body site and secondarily in each body site after bioinformatically pooling culture data from each atmosphere. To identify taxa that were differentially abundant in the hierarchical analysis, each taxonomic level from phylum to species was included for each individual ASV, when available. In assessing bacterial profile features preferentially recovered in one atmosphere over the other two, or in one body site over the other three, only mice with cultures from all three atmospheres in a body site were included (n = 9 for oral cavity, intestine, and vagina; n = 7 for the lung).

For all LEfSe analyses, singleton features were removed from each dataset, multi-class analysis of all-against-all was used only in identifying features that were preferentially abundant in one condition over all the others, and only features with an LDA score above 3.0 were considered preferentially abundant. Histograms (ASV-only analyses) and cladograms (hierarchical analyses) were generated using the Galaxy hub. Each taxon is indicated on cladograms when identified as a significant feature except order (to avoid visual congestion).

#### Mantel tests

Mantel tests were used to determine whether there was a correlation between the structure of bacterial culture profiles and the structure of molecular profiles for each body site. Only mice with bacterial profiles in both culture and molecular datasets in a body site were evaluated. Mantel tests were performed on Bray-Curtis distance matrices using the *vegan* package in RStudio (v 1.3.1093) and R (v. 4.0.3).

#### Figures

Heatmaps were generated using gplots and Heatplus packages in R (v. 4.0.3) and clustering of samples was performed on Bray-Curtis dissimilarity distance matrices using an unweighted pair group method with arithmetic mean (UPGMA) in the hclust function in R.

### Data availability

Original sample-specific MiSeq run files are available in the Short Read Archive from the original study Theis et al. (21), (BioProject identifier [ID] PRJNA594727). Assembled genomes of isolates ASV 2 and ASV 5 with annotations from NCBI’s Prokaryotic Genome Annotation Pipeline are available at BioProject ID PRJNA823350.

## REFERENCES

1. Krych L, Hansen CHF, Hansen AK, van den Berg FWJ, Nielsen DS. 2013. Quantitatively different, yet qualitatively alike: a meta-analysis of the mouse core gut microbiome with a view towards the human gut microbiome. PLOS ONE 8:e62578–e62578.

2. Walter J, Armet AM, Finlay BB, Shanahan F. 2020. Establishing or Exaggerating Causality for the Gut Microbiome: Lessons from Human Microbiota-Associated Rodents. Cell 180:221–232.

3. Nguyen TLA, Vieira-Silva S, Liston A, Raes J. 2015. How informative is the mouse for human gut microbiota research? Disease models & mechanisms 8:1–16.

4. Hugenholtz F, de Vos WM. 2018. Mouse models for human intestinal microbiota research: a critical evaluation. Cellular and molecular life sciences : CMLS 75:149–160.

5. Barfod KK, Roggenbuck M, Hansen LH, Schjørring S, Larsen ST, Sørensen SJ, Krogfelt KA. 2013. The murine lung microbiome in relation to the intestinal and vaginal bacterial communities. BMC microbiology 13:303–303.

6. Dollive S, Chen YY, Grunberg S, Bittinger K, Hoffmann C, Vandivier L, Bushman FD. 2013. Fungi of the murine gut: episodic variation and proliferation during antibiotic treatment. PLOS ONE 8.

7. Albenberg L, Esipova TV, Judge CP, Bittinger K, Chen J, Laughlin A, Grunberg S, Baldassano RN, Lewis JD, Li H, Thom SR, Bushman FD, Vinogradov SA, Wu GD. 2014. Correlation Between Intraluminal Oxygen Gradient and Radial Partitioning of Intestinal Microbiota. Gastroenterology 147:1055–1063.e8.

8. Braniste V, Al-Asmakh M, Kowal C, Anuar F, Abbaspour A, Tóth M, Korecka A, Bakocevic N, Ng LG, Kundu P, Gulyás B, Halldin C, Hultenby K, Nilsson H, Hebert H, Volpe BT, Diamond B, Pettersson S. 2014. The gut microbiota influences blood-brain barrier permeability in mice. Sci Transl Med 6:263ra158.

9. Koenigsknecht MJ, Theriot CM, Bergin IL, Schumacher CA, Schloss PD, Young VB. 2015. Dynamics and Establishment of Clostridium difficile Infection in the Murine Gastrointestinal Tract. Infection and Immunity 83:934–941.

10. Laukens D, Brinkman BM, Raes J, Vos M, Vandenabeele P. 2016. Heterogeneity of the gut microbiome in mice: guidelines for optimizing experimental design. FEMS Microbiol Rev 40.

11. Lagkouvardos I, Pukall R, Abt B, Foesel BU, Meier-Kolthoff JP, Kumar N, Bresciani A, Martínez I, Just S, Ziegler C, Brugiroux S, Garzetti D, Wenning M, Bui TPN, Wang J, Hugenholtz F, Plugge CM, Peterson DA, Hornef MW, Baines JF, Smidt H, Walter J, Kristiansen K, Nielsen HB, Haller D, Overmann J, Stecher B, Clavel T. 2016. The Mouse Intestinal Bacterial Collection (miBC) provides host-specific insight into cultured diversity and functional potential of the gut microbiota. Nature Microbiology 1:16131.

12. Tochitani S, Ikeno T, Ito T, Sakurai A, Yamauchi T, Matsuzaki H. 2016. Administration of Non-Absorbable Antibiotics to Pregnant Mice to Perturb the Maternal Gut Microbiota Is Associated with Alterations in Offspring Behavior. PLOS ONE 11:e0138293.

13. Elderman M, Hugenholtz F, Belzer C, Boekschoten M, de Haan B, de Vos P, Faas M. 2018. Changes in intestinal gene expression and microbiota composition during late pregnancy are mouse strain dependent. Scientific Reports 8:10001.

14. Wang J, Lang T, Shen J, Dai J, Tian L, Wang X. 2019. Core Gut Bacteria Analysis of Healthy Mice. Frontiers in Microbiology 10.

15. Vuong HE, Pronovost GN, Williams DW, Coley EJL, Siegler EL, Qiu A, Kazantsev M, Wilson CJ, Rendon T, Hsiao EY. 2020. The maternal microbiome modulates fetal neurodevelopment in mice. Nature 586:281–286.

16. Ahern PP, Maloy KJ. 2020. Understanding immune-microbiota interactions in the intestine. Immunology 159:4–14.

17. Kimura I, Miyamoto J, Ohue-Kitano R, Watanabe K, Yamada T, Onuki M, Aoki R, Isobe Y, Kashihara D, Inoue D, Inaba A, Takamura Y, Taira S, Kumaki S, Watanabe M, Ito M, Nakagawa F, Irie J, Kakuta H, Shinohara M, Iwatsuki K, Tsujimoto G, Ohno H, Arita M, Itoh H, Hase K. 2020. Maternal gut microbiota in pregnancy influences offspring metabolic phenotype in mice. Science 367:eaaw8429.

18. Karpinets TV, Solley TN, Mikkelson MD, Dorta-Estremera S, Nookala SS, Medrano AYD, Petrosino JF, Mezzari MP, Zhang J, Futreal PA, Sastry KJ, Colbert LE, Klopp A. 2020. Effect of Antibiotics on Gut and Vaginal Microbiomes Associated with Cervical Cancer Development in Mice. Cancer Prevention Research 13:997–1006.

19. Mandal RK, Denny JE, Waide ML, Li Q, Bhutiani N, Anderson CD, Baby BV, Jala VR, Egilmez NK, Schmidt NW. 2020. Temporospatial shifts within commercial laboratory mouse gut microbiota impact experimental reproducibility. BMC Biology 18:83.

20. Chung YW, Gwak H-J, Moon S, Rho M, Ryu J-H. 2020. Functional dynamics of bacterial species in the mouse gut microbiome revealed by metagenomic and metatranscriptomic analyses. PLOS ONE 15:e0227886.

21. Theis KR, Romero R, Greenberg JM, Winters AD, Garcia-Flores V, Motomura K, Ahmad MM, Galaz J, Arenas-Hernandez M, Gomez-Lopez N. 2020. No Consistent Evidence for Microbiota in Murine Placental and Fetal Tissues. mSphere 5:e00933–19.

22. Dudley DJ, Branch DW, Edwin SS, Mitchell MD. 1996. Induction of preterm birth in mice by RU486. Biol Reprod 55:992–5.

23. Hirsch E, Blanchard R, Mehta SP. 1999. Differential fetal and maternal contributions to the cytokine milieu in a murine model of infection-induced preterm birth. Am J Obstet Gynecol 180:429–34.

24. Elovitz MA, Mrinalini C. 2006. The use of progestational agents for preterm birth: lessons from a mouse model. Am J Obstet Gynecol 195:1004–10.

25. Shynlova O, Nedd-Roderique T, Li Y, Dorogin A, Lye SJ. 2013. Myometrial immune cells contribute to term parturition, preterm labour and post-partum involution in mice. J Cell Mol Med 17:90–102.

26. Akgul Y, Word RA, Ensign LM, Yamaguchi Y, Lydon J, Hanes J, Mahendroo M. 2014. Hyaluronan in cervical epithelia protects against infection-mediated preterm birth. J Clin Invest 124:5481–9.

27. Rinaldi SF, Makieva S, Frew L, Wade J, Thomson AJ, Moran CM, Norman JE, Stock SJ. 2015. Ultrasound-guided intrauterine injection of lipopolysaccharide as a novel model of preterm birth in the mouse. Am J Pathol 185:1201–6.

28. Vornhagen J, Quach P, Boldenow E, Merillat S, Whidbey C, Ngo LY, Adams Waldorf KM, Rajagopal L. 2016. Bacterial Hyaluronidase Promotes Ascending GBS Infection and Preterm Birth. mBio 7.

29. Gilman-Sachs A, Dambaeva S, Salazar Garcia MD, Hussein Y, Kwak-Kim J, Beaman K. 2018. Inflammation induced preterm labor and birth. J Reprod Immunol 129:53–58.

30. McCarthy R, Martin-Fairey C, Sojka DK, Herzog ED, Jungheim ES, Stout MJ, Fay JC, Mahendroo M, Reese J, Herington JL, Plosa EJ, Shelton EL, England SK. 2018. Mouse models of preterm birth: suggested assessment and reporting guidelines. Biol Reprod 99:922–937.

31. Gomez-Lopez N, Romero R, Arenas-Hernandez M, Panaitescu B, Garcia-Flores V, Mial TN, Sahi A, Hassan SS. 2018. Intra-amniotic administration of lipopolysaccharide induces spontaneous preterm labor and birth in the absence of a body temperature change. J Matern Fetal Neonatal Med 31:439–446.

32. Lee JY, Song H, Dash O, Park M, Shin NE, McLane MW, Lei J, Hwang JY, Burd I. 2019. Administration of melatonin for prevention of preterm birth and fetal brain injury associated with premature birth in a mouse model. Am J Reprod Immunol 82:e13151.

33. Nold C, Stone J, O’Hara K, Davis P, Kiveliyk V, Blanchard V, Yellon SM, Vella AT. 2019. Block of Granulocyte-Macrophage Colony-Stimulating Factor Prevents Inflammation-Induced Preterm Birth in a Mouse Model for Parturition. Reprod Sci 26:551–559.

34. Motomura K, Romero R, Xu Y, Theis KR, Galaz J, Winters AD, Slutsky R, Garcia-Flores V, Zou C, Levenson D, Para R, Ahmad MM, Miller D, Hsu CD, Gomez-Lopez N. 2020. Intra-Amniotic Infection with Ureaplasma parvum Causes Preterm Birth and Neonatal Mortality That Are Prevented by Treatment with Clarithromycin. mBio 11.

35. Gomez-Lopez N, Arenas-Hernandez M, Romero R, Miller D, Garcia-Flores V, Leng Y, Xu Y, Galaz J, Hassan SS, Hsu CD, Tse H, Sanchez-Torres C, Done B, Tarca AL. 2020. Regulatory T Cells Play a Role in a Subset of Idiopathic Preterm Labor/Birth and Adverse Neonatal Outcomes. Cell Rep 32:107874.

36. Fan C, Dai Y, Zhang L, Rui C, Wang X, Luan T, Fan Y, Dong Z, Hou W, Li P, Liao Q, Zeng X. 2021. Aerobic Vaginitis Induced by Escherichia coli Infection During Pregnancy Can Result in Adverse Pregnancy Outcomes Through the IL-4/JAK-1/STAT-6 Pathway. Front Microbiol 12:651426.

37. Spencer NR, Radnaa E, Baljinnyam T, Kechichian T, Tantengco OAG, Bonney E, Kammala AK, Sheller-Miller S, Menon R. 2021. Development of a mouse model of ascending infection and preterm birth. PLOS ONE 16:e0260370.

38. Gershater M, Romero R, Arenas-Hernandez M, Galaz J, Motomura K, Tao L, Xu Y, Miller D, Pique-Regi R, Martinez G, 3rd, Liu Y, Jung E, Para R, Gomez-Lopez N. 2022. IL-22 Plays a Dual Role in the Amniotic Cavity: Tissue Injury and Host Defense against Microbes in Preterm Labor. J Immunol 208:1595–1615.

39. Faas MM, Liu Y, Borghuis T, van Loo-Bouwman CA, Harmsen H, de Vos P. 2019. Microbiota Induced Changes in the Immune Response in Pregnant Mice. Front Immunol 10:2976.

40. Gravett MG, Hummel D, Eschenbach DA, Holmes KK. 1986. Preterm labor associated with subclinical amniotic fluid infection and with bacterial vaginosis. Obstet Gynecol 67:229–37.

41. Krohn MA, Hillier SL, Nugent RP, Cotch MF, Carey JC, Gibbs RS, Eschenbach DA. 1995. The genital flora of women with intraamniotic infection. Vaginal Infection and Prematurity Study Group. J Infect Dis 171:1475–80.

42. Romero R, Gomez-Lopez N, Winters AD, Jung E, Shaman M, Bieda J, Panaitescu B, Pacora P, Erez O, Greenberg JM, Ahmad MM, Hsu CD, Theis KR. 2019. Evidence that intra-amniotic infections are often the result of an ascending invasion - a molecular microbiological study. J Perinat Med 47:915–931.

43. Cobo T, Vergara A, Collado MC, Casals-Pascual C, Herreros E, Bosch J, Sánchez-García AB, López-Parellada R, Ponce J, Gratacós E. 2019. Characterization of vaginal microbiota in women with preterm labor with intra-amniotic inflammation. Sci Rep 9:18963.

44. Eschenbach DA, Gravett MG, Chen KC, Hoyme UB, Holmes KK. 1984. Bacterial vaginosis during pregnancy. An association with prematurity and postpartum complications. Scand J Urol Nephrol Suppl 86:213–22.

45. Holst E, Goffeng AR, Andersch B. 1994. Bacterial vaginosis and vaginal microorganisms in idiopathic premature labor and association with pregnancy outcome. J Clin Microbiol 32:176–86.

46. Hillier SL, Nugent RP, Eschenbach DA, Krohn MA, Gibbs RS, Martin DH, Cotch MF, Edelman R, Pastorek JG, 2nd, Rao AV, et al. 1995. Association between bacterial vaginosis and preterm delivery of a low-birth-weight infant. The Vaginal Infections and Prematurity Study Group. N Engl J Med 333:1737–42.

47. Meis PJ, Goldenberg RL, Mercer B, Moawad A, Das A, McNellis D, Johnson F, Iams JD, Thom E, Andrews WW. 1995. The preterm prediction study: significance of vaginal infections. National Institute of Child Health and Human Development Maternal-Fetal Medicine Units Network. Am J Obstet Gynecol 173:1231–5.

48. Nelson DB, Bellamy S, Nachamkin I, Ness RB, Macones GA, Allen-Taylor L. 2007. First trimester bacterial vaginosis, individual microorganism levels, and risk of second trimester pregnancy loss among urban women. Fertility and Sterility 88:1396–1403.

49. Giraldo PC, Araujo ED, Junior JE, do Amaral RL, Passos MR, Goncalves AK. 2012. The prevalence of urogenital infections in pregnant women experiencing preterm and full-term labor. Infect Dis Obstet Gynecol 2012:878241.

50. Romero R, Hassan SS, Gajer P, Tarca AL, Fadrosh DW, Bieda J, Chaemsaithong P, Miranda J, Chaiworapongsa T, Ravel J. 2014. The vaginal microbiota of pregnant women who subsequently have spontaneous preterm labor and delivery and those with a normal delivery at term. Microbiome 2:18.

51. Digiulio DB, Callahan BJ, McMurdie PJ, Costello EK, Lyell DJ, Robaczewska A, Sun CL, Goltsman DSA, Wong RJ, Shaw G, Stevenson DK, Holmes SP, Relman DA. 2015. Temporal and spatial variation of the human microbiota during pregnancy. Proceedings of the National Academy of Sciences 112:11060–11065.

52. Cox C, Saxena N, Watt AP, Gannon C, McKenna JP, Fairley DJ, Sweet D, Shields MD, Cosby SL, Coyle PV. 2016. The common vaginal commensal bacterium Ureaplasma parvum is associated with chorioamnionitis in extreme preterm labor. Journal of Maternal-Fetal & Neonatal Medicine 29:3646–3651.

53. Haque MM, Merchant M, Kumar PN, Dutta A, Mande SS. 2017. First-trimester vaginal microbiome diversity: A potential indicator of preterm delivery risk. Scientific Reports 7.

54. Elovitz MA, Gajer P, Riis V, Brown AG, Humphrys MS, Holm JB, Ravel J. 2019. Cervicovaginal microbiota and local immune response modulate the risk of spontaneous preterm delivery. Nat Commun 10:1305.

55. Fettweis JM, Serrano MG, Brooks JP, Edwards DJ, Girerd PH, Parikh HI, Huang B, Arodz TJ, Edupuganti L, Glascock AL, Xu J, Jimenez NR, Vivadelli SC, Fong SS, Sheth NU, Jean S, Lee V, Bokhari YA, Lara AM, Mistry SD, Duckworth RA, 3rd, Bradley SP, Koparde VN, Orenda XV, Milton SH, Rozycki SK, Matveyev AV, Wright ML, Huzurbazar SV, Jackson EM, Smirnova E, Korlach J, Tsai YC, Dickinson MR, Brooks JL, Drake JI, Chaffin DO, Sexton AL, Gravett MG, Rubens CE, Wijesooriya NR, Hendricks-Muñoz KD, Jefferson KK, Strauss JF, 3rd, Buck GA. 2019. The vaginal microbiome and preterm birth. Nat Med 25:1012–1021.

56. Zhang F, Zhang T, Ma Y, Huang Z, He Y, Pan H, Fang M, Ding H. 2019. Alteration of vaginal microbiota in patients with unexplained recurrent miscarriage. Experimental and therapeutic medicine 17:3307–3316.

57. Al-Memar M, Bobdiwala S, Fourie H, Mannino R, Lee Y, Smith A, Marchesi J, Timmerman D, Bourne T, Bennett P, MacIntyre D. 2020. The association between vaginal bacterial composition and miscarriage: a nested case–control study. BJOG: An International Journal of Obstetrics & Gynaecology 127:264–274.

58. Chan D, Bennett PR, Lee YS, Kundu S, Teoh TG, Adan M, Ahmed S, Brown RG, David AL, Lewis HV, Gimeno-Molina B, Norman JE, Stock SJ, Terzidou V, Kropf P, Botto M, MacIntyre DA, Sykes L. 2022. Microbial-driven preterm labour involves crosstalk between the innate and adaptive immune response. Nat Commun 13:975.

59. Miller EA, Beasley DE, Dunn RR, Archie EA. 2016. Lactobacilli Dominance and Vaginal pH: Why Is the Human Vaginal Microbiome Unique? Front Microbiol 7:1936.

60. Amabebe E, Anumba DOC. 2018. The Vaginal Microenvironment: The Physiologic Role of Lactobacilli. Front Med (Lausanne) 5:181.

61. Saraf VS, Sheikh SA, Ahmad A, Gillevet PM, Bokhari H, Javed S. 2021. Vaginal microbiome: normalcy vs dysbiosis. Arch Microbiol doi:10.1007/s00203-021-02414-3.

62. Ravel J, Gajer P, Abdo Z, Schneider GM, Koenig SS, McCulle SL, Karlebach S, Gorle R, Russell J, Tacket CO, Brotman RM, Davis CC, Ault K, Peralta L, Forney LJ. 2011. Vaginal microbiome of reproductive-age women. Proc Natl Acad Sci U S A 108 Suppl 1:4680–7.

63. Chang DH, Shin J, Rhee MS, Park KR, Cho BK, Lee SK, Kim BC. 2020. Vaginal Microbiota Profiles of Native Korean Women and Associations with High-Risk Pregnancy. J Microbiol Biotechnol 30:248–258.

64. Dunlop AL, Satten GA, Hu YJ, Knight AK, Hill CC, Wright ML, Smith AK, Read TD, Pearce BD, Corwin EJ. 2021. Vaginal Microbiome Composition in Early Pregnancy and Risk of Spontaneous Preterm and Early Term Birth Among African American Women. Front Cell Infect Microbiol 11:641005.

65. Kumar S, Kumari N, Talukdar D, Kothidar A, Sarkar M, Mehta O, Kshetrapal P, Wadhwa N, Thiruvengadam R, Desiraju BK, Nair GB, Bhatnagar S, Mukherjee S, Das B. 2021. The Vaginal Microbial Signatures of Preterm Birth Delivery in Indian Women. Front Cell Infect Microbiol 11:622474.

66. Romero R, Hassan SS, Gajer P, Tarca AL, Fadrosh DW, Nikita L, Galuppi M, Lamont RF, Chaemsaithong P, Miranda J, Chaiworapongsa T, Ravel J. 2014. The composition and stability of the vaginal microbiota of normal pregnant women is different from that of non-pregnant women. Microbiome 2:4.

67. Han YW, Ikegami A, Bissada NF, Herbst M, Redline RW, Ashmead GG. 2006. Transmission of an uncultivated Bergeyella strain from the oral cavity to amniotic fluid in a case of preterm birth. J Clin Microbiol 44:1475–83.

68. Han YW, Fardini Y, Chen C, Iacampo KG, Peraino VA, Shamonki JM, Redline RW. 2010. Term stillbirth caused by oral Fusobacterium nucleatum. Obstet Gynecol 115:442–5.

69. Davis EP. 1890. TUBERCULOSIS IN PREGNANCY AND PARTURITION.-- WEBBED FINGER.--DIFFERENTIAL DIAGNOSIS OF PNEUMONIA IN CHILDREN.: TUBERCULOSIS IN PREGNANCY AND PARTURITION. WEBBED FINGERS. CATARRHAL AND CROUPOUS PNEUMONIA. Medical and Surgical Reporter (1858-1898) 62:420.

70. Huber BM, Meyer Sauteur PM, Unger WWJ, Hasters P, Eugster MR, Brandt S, Bloemberg GV, Natalucci G, Berger C. 2018. Vertical Transmission of Mycoplasma pneumoniae Infection. Neonatology 114:332–336.

71. Winn HN. 2007. Group B Streptococcus Infection in Pregnancy. Clinics in Perinatology 34:387–392.

72. Larsen JW, Sever JL. 2008. Group B Streptococcus and pregnancy: a review. American Journal of Obstetrics and Gynecology 198:440–450.

73. Brokaw A, Furuta A, Dacanay M, Rajagopal L, Adams Waldorf KM. 2021. Bacterial and Host Determinants of Group B Streptococcal Vaginal Colonization and Ascending Infection in Pregnancy. Front Cell Infect Microbiol 11:720789.

74. Waite DW, Chuvochina M, Pelikan C, Parks DH, Yilmaz P, Wagner M, Loy A, Naganuma T, Nakai R, Whitman WB, Hahn MW, Kuever J, Hugenholtz P. 2020. Proposal to reclassify the proteobacterial classes Deltaproteobacteria and Oligoflexia, and the phylum Thermodesulfobacteria into four phyla reflecting major functional capabilities. International Journal of Systematic and Evolutionary Microbiology 70:5972–6016.

75. Kent WJ. 2002. BLAT - The BLAST-like alignment tool. Genome Research 12:656–664.

76. Kanehisa M, Sato Y. 2020. KEGG Mapper for inferring cellular functions from protein sequences. Protein Science 29:28–35.

77. Seemann T. 2014. Prokka: rapid prokaryotic genome annotation. Bioinformatics 30:2068–9.

78. Madigan MT, Martinko JM, Parker J. 2006. Brock biology of microorganisms, vol 11. Pearson Prentice Hall Upper Saddle River, NJ.

79. Morris RL, Schmidt TM. 2013. Shallow breathing: bacterial life at low O2. Nature Reviews Microbiology 11:205–212.

80. Abusleme L, O’Gorman H, Dutzan N, Greenwell-Wild T, Moutsopoulos NM. 2020. Establishment and Stability of the Murine Oral Microbiome. Journal of Dental Research 99:721–729.

81. Rautava J, Pinnell LJ, Vong L, Akseer N, Assa A, Sherman PM. 2015. Oral microbiome composition changes in mouse models of colitis. Journal of Gastroenterology and Hepatology 30:521–527.

82. Abusleme L, Hong B-Y, Hoare A, Konkel JE, Diaz PI, Moutsopoulos NM. 2017. Oral Microbiome Characterization in Murine Models. Bio-protocol 7:e2655.

83. A. Hernández-Arriaga A, Baumann A, Witte OW, Frahm C, Bergheim I, Camarinha-Silva A. 2019. Changes in Oral Microbial Ecology of C57BL/6 Mice at Different Ages Associated with Sampling Methodology. Microorganisms 7:283.

84. Dickson RP, Erb-Downward JR, Falkowski NR, Hunter EM, Ashley SL, Huffnagle GB. 2018. The Lung Microbiota of Healthy Mice Are Highly Variable, Cluster by Environment, and Reflect Variation in Baseline Lung Innate Immunity. American journal of respiratory and critical care medicine 198:497–508.

85. Singh N, Vats A, Sharma A, Arora A, Kumar A. 2017. The development of lower respiratory tract microbiome in mice. Microbiome 5:61–61.

86. A. Kostric M, Milger K, Krauss-Etschmann S, Engel M, Vestergaard G, Schloter M, Schöler A. 2018. Development of a Stable Lung Microbiome in Healthy Neonatal Mice. Microbial Ecology 75:529–542.

87. Ashley SL, Sjoding MW, Popova AP, Cui TX, Hoostal MJ, Schmidt TM, Branton WR, Dieterle MG, Falkowski NR, Baker JM, Hinkle KJ, Konopka KE, Erb-Downward JR, Huffnagle GB, Dickson RP. 2020. Lung and gut microbiota are altered by hyperoxia and contribute to oxygen-induced lung injury in mice. Science Translational Medicine 12:eaau9959.

88. Poroyko V, Meng F, Meliton A, Afonyushkin T, Ulanov A, Semenyuk E, Latif O, Tesic V, Birukova AA, Birukov KG. 2015. Alterations of lung microbiota in a mouse model of LPS-induced lung injury. American journal of physiology Lung cellular and molecular physiology 309:L76–L83.

89. Vrbanac A, Riestra AM, Coady A, Knight R, Nizet V, Patras KA. 2018. The murine vaginal microbiota and its perturbation by the human pathogen group B Streptococcus. BMC Microbiology 18:197.

90. Cowley HM, Heiss GS. 1991. Changes in Vaginal Bacterial Flora During the Oestrous Cycle of the Mouse. Microbial Ecology in Health and Disease 4:229–235.

91. Noguchi K, Tsukumi K, Urano T. 2003. Qualitative and Quantitative Differences in Normal Vaginal Flora of Conventionally Reared Mice, Rats, Hamsters, Rabbits, and Dogs. Comparative Medicine 53:404–412.

92. Jašarević E, Howard CD, Misic AM, Beiting DP, Bale TL. 2017. Stress during pregnancy alters temporal and spatial dynamics of the maternal and offspring microbiome in a sex-specific manner. Scientific Reports 7:44182.

93. Younge N, McCann JR, Ballard J, Plunkett C, Akhtar S, Araújo-Pérez F, Murtha A, Brandon D, Seed PC. 2019. Fetal exposure to the maternal microbiota in humans and mice. JCI Insight 4.

94. Adhikary S, Nicklas W, Bisgaard M, Boot R, Kuhnert P, Waberschek T, Aalbæk B, Korczak B, Christensen H. 2017. Rodentibacter gen. nov. including Rodentibacter pneumotropicus comb. nov., Rodentibacter heylii sp. nov., Rodentibacter myodis sp. nov., Rodentibacter ratti sp. nov., Rodentibacter heidelbergensis sp. nov., Rodentibacter trehalosifermentans sp. nov., Rodentibacter rarus sp. nov., Rodentibacter mrazii and two genomospecies. International Journal of Systematic and Evolutionary Microbiology 67:1793–1806.

95. Biggers JD. 1953. The carbohydrate components of the vagina of the normal and ovariectomized mouse during oestrogenic stimulation. Journal of anatomy 87:327–336.

96. Dal Bello F, Hertel C. 2006. Oral cavity as natural reservoir for intestinal lactobacilli. Systematic and Applied Microbiology 29:69–76.

97. Ursell LK, Clemente JC, Rideout JR, Gevers D, Caporaso JG, Knight R. 2012. The interpersonal and intrapersonal diversity of human-associated microbiota in key body sites. Journal of Allergy and Clinical Immunology 129:1204–1208.

98. Arumugam M, Raes J, Pelletier E, Le Paslier D, Yamada T, Mende DR, Fernandes GR, Tap J, Bruls T, Batto JMM, Bertalan M, Borruel N, Casellas F, Fernandez L, Gautier L, Hansen T, Hattori M, Hayashi T, Kleerebezem M, Kurokawa K, Leclerc M, Levenez F, Manichanh C, Nielsen HB, Nielsen T, Pons N, Poulain J, Qin J, Sicheritz-Ponten T, Tims S. 2011. Enterotypes of the human gut microbiome. Nature 473.

99. Lozupone CA, Stombaugh JI, Gordon JI, Jansson JK, Knight R. 2012. Diversity, stability and resilience of the human gut microbiota. Nature 489.

100. Lloyd-Price J, Mahurkar A, Rahnavard G, Crabtree J, Orvis J, Hall AB, Brady A, Creasy HH, McCracken C, Giglio MG, McDonald D, Franzosa EA, Knight R, White O, Huttenhower C. 2017. Strains, functions and dynamics in the expanded Human Microbiome Project. Nature 550:61–66.

101. Benga L, Knorr JI, Engelhardt E, Gougoula C, Benten PM, Christensen H, Sager M. 2019. Current Distribution of Rodentibacter Species Among the Mice and Rats of an Experimental Facility. Journal of the American Association for Laboratory Animal Science 58:475–478.

102. Mirmonsef P, Hotton AL, Gilbert D, Burgad D, Landay A, Weber KM, Cohen M, Ravel J, Spear GT. 2014. Free glycogen in vaginal fluids is associated with Lactobacillus colonization and low vaginal pH. PLOS ONE 9:e102467–e102467.

103. Rausch P, Basic M, Batra A, Bischoff SC, Blaut M, Clavel T, Gläsner J, Gopalakrishnan S, Grassl GA, Günther C, Haller D, Hirose M, Ibrahim S, Loh G, Mattner J, Nagel S, Pabst O, Schmidt F, Siegmund B, Strowig T, Volynets V, Wirtz S, Zeissig S, Zeissig Y, Bleich A, Baines JF. 2016. Analysis of factors contributing to variation in the C57BL/6J fecal microbiota across German animal facilities. International Journal of Medical Microbiology 306:343–355.

104. Witjes VM, Boleij A, Halffman W. 2020. Reducing versus Embracing Variation as Strategies for Reproducibility: The Microbiome of Laboratory Mice. Animals 10:2415.

105. Stevenson BS, Eichorst SA, Wertz JT, Schmidt TM, Breznak JA. 2004. New strategies for cultivation and detection of previously uncultured microbes. Appl Environ Microbiol 70:4748–55.

106. Kozich JJ, Westcott SL, Baxter NT, Highlander SK, Schloss PD. 2013. Development of a dual-index sequencing strategy and curation pipeline for analyzing amplicon sequence data on the MiSeq Illumina sequencing platform. Appl Environ Microbiol 79:5112–20.

107. Callahan BJ, McMurdie PJ, Rosen MJ, Han AW, Johnson AJA, Holmes SP. 2016. DADA2: High-resolution sample inference from Illumina amplicon data. Nature Methods 13:581–583.

108. Pruesse E, Quast C, Knittel K, Fuchs BM, Ludwig W, Peplies J, Glockner FO. 2007. SILVA: a comprehensive online resource for quality checked and aligned ribosomal RNA sequence data compatible with ARB. Nucleic Acids Research 35:7188–7196.

109. Quast C, Pruesse E, Yilmaz P, Gerken J, Schweer T, Yarza P, Peplies J, Glockner FO. 2013. The SILVA ribosomal RNA gene database project: improved data processing and web-based tools. Nucleic Acids Res 41:D590–6.

110. Davis NM, Proctor DM, Holmes SP, Relman DA, Callahan BJ. 2018. Simple statistical identification and removal of contaminant sequences in marker-gene and metagenomics data. Microbiome 6:226.

111. Afgan E, Baker D, Batut B, van den Beek M, Bouvier D, Čech M, Chilton J, Clements D, Coraor N, Grüning BA, Guerler A, Hillman-Jackson J, Hiltemann S, Jalili V, Rasche H, Soranzo N, Goecks J, Taylor J, Nekrutenko A, Blankenberg D. 2018. The Galaxy platform for accessible, reproducible and collaborative biomedical analyses: 2018 update. Nucleic Acids Research 46:W537–W544.

112. Na S-I, Kim YO, Yoon S-H, Ha S-m, Baek I, Chun J. 2018. UBCG: Up-to-date bacterial core gene set and pipeline for phylogenomic tree reconstruction. Journal of Microbiology 56:280–285.

113. Rambaut A. 2018. FigTree v1.4.4, Institute of Evolutionary Biology, University of Edinburgh, Edinburgh. https://github.com/rambaut/figtree.

114. Darling ACE, Mau B, Blattner FR, Perna NT. 2004. Mauve: multiple alignment of conserved genomic sequence with rearrangements. Genome research 14:1394–1403.

115. Page AJ, Cummins CA, Hunt M, Wong VK, Reuter S, Holden MTG, Fookes M, Falush D, Keane JA, Parkhill J. 2015. Roary: rapid large-scale prokaryote pan genome analysis. Bioinformatics 31:3691–3693.

116. Tatusova T, DiCuccio M, Badretdin A, Chetvernin V, Nawrocki EP, Zaslavsky L, Lomsadze A, Pruitt KD, Borodovsky M, Ostell J. 2016. NCBI prokaryotic genome annotation pipeline. Nucleic Acids Research 44:6614–6624.

117. Moriya Y, Itoh M, Okuda S, Yoshizawa AC, Kanehisa M. 2007. KAAS: an automatic genome annotation and pathway reconstruction server. Nucleic Acids Res 35:W182–5.

118. Karp PD, Midford PE, Billington R, Kothari A, Krummenacker M, Latendresse M, Ong WK, Subhraveti P, Caspi R, Fulcher C, Keseler IM, Paley SM. 2021. Pathway Tools version 23.0 update: software for pathway/genome informatics and systems biology. Brief Bioinform 22:109–126.

119. Gohir W, Whelan FJ, Surette MG, Moore C, Schertzer JD, Sloboda DM. 2015. Pregnancy-related changes in the maternal gut microbiota are dependent upon the mother’s periconceptional diet. Gut Microbes 6:310–320.

120. Nuriel-Ohayon M, Neuman H, Ziv O, Belogolovski A, Barsheshet Y, Bloch N, Uzan A, Lahav R, Peretz A, Frishman S, Hod M, Hadar E, Louzoun Y, Avni O, Koren O. 2019. Progesterone Increases Bifidobacterium Relative Abundance during Late Pregnancy. Cell Reports 27:730–736.e3.

121. Liu X, Zhang F, Wang Z, Zhang T, Teng C, Wang Z. 2021. Altered gut microbiome accompanying with placenta barrier dysfunction programs pregnant complications in mice caused by graphene oxide. Ecotoxicology and Environmental Safety 207:111143.

