## Supplemental Materials for "Microbiota of the pregnant mouse: characterization of the bacterial communities in the oral cavity, lung, intestine, and vagina through culture and DNA sequencing"

### CORRESPONDING AUTHOR(S)

**Supplemental Table 1.** Description of previous 16S rRNA gene studies of the murine oral microbiota.

| Author(s) | Year | Culture methods | Mouse strain | Mouse sex | Key microbiota findings |
|---|---|---|---|---|---|
| Abusleme, et al. (1) | 2017 | Not performed | C57BL/6 | Not indicated | The most abundant genera observed in the oral cavity were *Lactobacillus, Staphyloccocus, Enterococcus, Rothia, Acinetobacter*, and *Streptococcus.* |
| Hernández-Arriaga, et al. (2) | 2019 | Not performed | C57BL/6 | Exclusively male | Relatively abundant genera varied with age and sampling method. Relatively abundant genera included *Actinobacillus, Actinomyces, Neisseria, Corynebacterium, Aggregatibacter, Staphylococcus, Streptococcus*, and *Propionibacterium. Rodentibacter* was observed in some mice. |
| Abusleme, et al. (3) | 2020 | Not performed | C57BL/6 | Not indicated | Bacteria relatively abundant in adult mice included *Streptococcus, Lactobacillus* spp., *Turicibacter, Staphylococcus, Enterococcus,* and *Flavobacterium.* Bacterial composition was distinct for mice from different laboratories. |
| Jašarević, et al. (4) | 2021 | Not performed | C57BL/6 | Exclusively female | Prominent taxa were Lachnospiraceae, Bacteroidales, *Oscillibacter*, and *Draconibacterium.* |

**Supplemental Table 2.** Description of previous 16S rRNA gene studies of the murine lung microbiota.

| Author(s) | Year | Culture methods | Mouse strain | Mouse sex | Key microbiota findings |
|---|---|---|---|---|---|
| Barfod, et al. (5) | 2013 | 200uL of bronchoalveolar lavage (BAL) fluid was cultured on general growth media blood agar 5% and Chocolate agar at 37ºC for 24 hours and under hypoxic conditions (5% $CO_2$, 3% $H_2$, 5% $O_2$ and 87% $N_2$) for 48 hours and identified by a Vitek2 system | BALB/cJ | Exclusively female | 12 core genera were observed between lung, cecum, and vaginal samples. The lung and vagina shared many genera. *Staphylococcus*, *Massilia*, *Corynebacterium*, *Pseudomonas*, *Streptococcus*, and *Sphingomonas* were detected in all lung samples. Only *Micrococcus luteus* was recovered via culture. |
| Poroyko, et al. (6) | 2015 | A single drop of BAL fluid was added to chopped meat carbohydrate broth, PR II, tryptic soy agar with 5% sheep blood, and chocolate II agar plates and incubated at 5% $CO_2$ at 37ºC and Brucella agar with 5% sheep blood, hemin, and vitamin K1 incubated under anaerobic conditions. Plates were checked daily for 4 days and broth was checked daily for 14 days. | C57BL/6 | Not mentioned | Molecular surveys detected a high relative abundance of the family Alicyclobacillaceae. *Stenotrophomonas* and *Ochrobactrum* were observed in culture and molecular surveys. |
| Singh, et al. (7) | 2017 | Not performed | C57BL/6N | Yes, but not addressed in analyses. | Abundant genera observed in adult mice included *Actinobacillus*, *Achromobacter*, *Lactobacillus*, *Bacillus*, and *Streptococcus*. |
| Dickson, et al. (8) | 2018 | Not performed | C57BL/6 | Exclusively female | No core lung microbiome was observed. Environmental factors such as cohabitation and supplying vendor were primary drivers in microbiome composition. Most abundant taxa observed include *Pseudomonas*, *Lactobacillus*, |

| | | | | | |
|---|---|---|---|---|---|
| | | | | | *Streptococcus* and *Turicibacter*. |
| Kostric, et al. (9) | 2018 | Not performed | BALB/c | Exclusively female | Abundant genera in adult (2-9 mo) mice included *Rhodococcus*, *Staphylococcus*, *Lactobacillus*, *Streptococcus*, *Delftia*, *Propionibacterium*, *Pseudomonas*, *Diaphorobacter*, *Defluvibacter*, *Ochrobactrum*, *Mucispirillium*. |
| Ashley, et al. (10) | 2020 | Not performed | C57BL/6 | Exclusively female | Abundant genera observed in control mice included *Pseudomonas*, *Lactobacillus*, and *Bacteroides*. |
| Li, et al. (11) | 2020 | Not performed | C57BL/6N | No | Microbiota characterization was limited to phyla. |
| Baker, et al. (12) | 2021 | Not performed | C57BL/6 | Exclusively female | Top taxa were *Turicibacter*, *Lactobacillus*, *Flavobacterium*, and *Alistipes*. |
| Lipinski, et al. (13) | 2021 | Not performed | C57BL/6 | Exclusively female | *Sphingomonas* and *Burkholderia* were the most relatively abundant taxa among control mice. |

**Supplemental Table 3.** Description of previous 16S rRNA gene studies of the murine vaginal microbiota.

| Author(s) | Year | Culture methods | Mouse strain | Mouse sex | Key microbiota findings |
|---|---|---|---|---|---|
| Barfod, et al. (5) | 2013 | Not performed | BALB/cJ | N/A | Vaginal samples from mice clustered into two groups and were typically abundant in *Streptococcus.* Other notable genera included *Acinetobacter, Sphingomonas, Enterococcus*, and *Polaromonas*. |
| Oh, et al. (14) | 2016 | Vaginal wash fluids were spread onto CNA and MacConkey agar plates at 35ºC for 48h under oxic conditions and on anaerobic basal agar Oxoid plates at 37ºC for 5 days under anaerobic conditions. | C57BL/6 | N/A | Vaginal washes from control mice were composed of AY538685, *Proteus*, *Escherichia*, *Enterobacter*, *Serratia*, and *Staphylococcus*. Culture was used strictly to assess bacterial load. |
| Vrbanac, et al. (15) | 2018 | Not performed | C57BL/6J | N/A | Five distinctive community states of the vaginal microbiota were described. These community states were defined by abundance in *Staphylococcus* and/or *Enterococcus*, *Lactobacillus*, or a mixed population. The stage of estrous did not impact vaginal community structure. |
| Karpinets, et al. (16) | 2020 | Not performed | C57BL/6 | N/A | Relatively abundant genera found in control mice included *Bacteroides, Staphylococcus, Turicibacter, Streptococcus, Paraliobacillus, Lachnoclostridium, Paeniclostridium, Halorubellus, Halorubrum, Lactobacillus, Oscillibacter, Romboutsia, Parabacteroides, Atopostipes, Akkermansia, and Sporosarcina.* |
| Hernandez-Quiroz, et al. (17) | 2021 | Not performed | FVB | N/A | Relatively abundant taxa included *Streptococcus*, *Microbacterium, Kaistobacter*, *Aggregatibacter*, and Weeksellaceae. |
| Jašarević, et al. (4) | 2021 | Not performed | C57BL/6 | N/A | The vagina was most relatively abundant in *Pasteurella* |

| | | | | | |
|---|---|---|---|---|---|
| | | | | | *pneumotropica* (recently reclassified as *Rodentibacter*). |
| Mejia, et al. (18) | 2022 | Not performed | C57BL/6J | N/A | Similar to Vrbanac, et al., microbiota were classified into CSTs according to the dominant taxa *Staphylococcus*, *Enterococcus*, or *Lactobacillus*. |

**Supplemental Table 4.** Comparisons of the bacterial profiles of cultured murine sample microbiota under anoxic, hypoxic, and oxic conditions.

| **Beta diversity** | **Composition** | | | **Structure** | | |
|---|---|---|---|---|---|---|
| | **F** | **$R^2$** | **p** | **F** | **$R^2$** | **p** |
| **Anoxic** | | | | | | |
| Mouse ID | 1.049 | 0.224 | 0.299 | 1.107 | 0.207 | 0.252 |
| Body site | 3.745 | 0.240 | **0.001** | 5.830 | 0.327 | **0.001** |
| **Hypoxic** | | | | | | |
| Mouse ID | 1.190 | 0.268 | **0.019** | 0.766 | 0.195 | 0.923 |
| Body site | 2.845 | 0.192 | **0.001** | 2.548 | 0.194 | **0.001** |
| **Oxic** | | | | | | |
| Mouse ID | 1.089 | 0.249 | 0.175 | 1.307 | 0.309 | 0.101 |
| Body site | 2.936 | 0.202 | **0.001** | 1.764 | 0.125 | **0.039** |

**Supplemental Table 5.** Comparisons of the (A) composition and (B) structure of the cultured microbiota between different atmospheres for the oral cavity, lung, intestine, and vagina.

| **A** | **Composition** | | | | | | | | | | | |
|---|---|---|---|---|---|---|---|---|---|---|---|---|
| **Bacterial composition** | **All atmospheres** | | | **Oxic x Hypoxic** | | | **Oxic x Anoxic** | | | **Hypoxic x Anoxic** | | |
| | **F** | **$R^2$** | **p** | **F** | **$R^2$** | **p** | **F** | **$R^2$** | **p** | **F** | **$R^2$** | **p** |
| **Oral cavity** | | | | | | | | | | | | |
| Mouse ID | 0.981 | 0.327 | 0.569 | 1.201 | 0.535 | 0.089 | 0.784 | 0.448 | 0.970 | 0.980 | 0.481 | 0.544 |
| Atmosphere | 1.581 | 0.106 | **0.015** | 1.387 | 0.069 | 0.111 | 1.659 | 0.095 | **0.040** | 1.583 | 0.078 | **0.042** |
| **Lung** | | | | | | | | | | | | |
| Mouse ID | 1.039 | 0.348 | 0.316 | 0.904 | 0.507 | 0.841 | 1.050 | 0.515 | 0.307 | 1.147 | 0.565 | 0.131 |
| Atmosphere | 1.268 | 0.094 | 0.070 | | | | | | | | | |
| **Intestine** | | | | | | | | | | | | |
| Mouse ID | 1.029 | 0.280 | 0.381 | 0.974 | 0.463 | 0.570 | 1.016 | 0.412 | 0.437 | 1.127 | 0.449 | 0.205 |
| Atmosphere | 3.435 | 0.207 | **0.001** | 1.150 | 0.061 | 0.264 | 5.038 | 0.227 | **0.001** | 4.456 | 0.197 | **0.001** |
| **Vagina** | | | | | | | | | | | | |
| Mouse ID[†] | 1.911 | 0.511 | **0.001** | | | | | | | | | |
| Atmosphere[‡] | 1.142 | 0.061 | 0.202 | 0.682 | 0.039 | 0.566 | 1.059 | 0.056 | 0.084 | 0.938 | 0.052 | 0.063 |
| **B** | **Structure** | | | | | | | | | | | |
| **Bacterial structure** | **All atmospheres** | | | **Oxic x Hypoxic** | | | **Oxic x Anoxic** | | | **Hypoxic x Anoxic** | | |
| | **F** | **$R^2$** | **p** | **F** | **$R^2$** | **p** | **F** | **$R^2$** | **p** | **F** | **$R^2$** | **p** |
| **Oral cavity** | | | | | | | | | | | | |
| Mouse ID | 1.005 | 0.296 | 0.494 | 2.794 | 0.719 | **0.013*** | 0.706 | 0.359 | 0.861 | 0.617 | 0.343 | 0.956 |
| Atmosphere | 3.434 | 0.202 | **0.007** | 1.832 | 0.052 | 0.138 | 4.597 | 0.234 | **0.004** | 2.836 | 0.157 | **0.026** |
| **Lung** | | | | | | | | | | | | |
| Mouse ID | 0.847 | 0.302 | 0.774 | 0.801 | 0.481 | 0.811 | 0.931 | 0.490 | 0.621 | 0.886 | 0.502 | 0.708 |
| Atmosphere | 1.313 | 0.104 | 0.167 | | | | | | | | | |
| **Intestine** | | | | | | | | | | | | |
| Mouse ID | 1.205 | 0.300 | 0.202 | 1.058 | 0.488 | 0.363 | 1.072 | 0.384 | 0.371 | 1.537 | 0.506 | 0.097 |
| Atmosphere | 4.161 | 0.230 | **0.001** | 0.969 | 0.050 | 0.425 | 7.464 | 0.297 | **0.001** | 5.528 | 0.202 | **0.001** |
| **Vagina** | | | | | | | | | | | | |
| Mouse ID[†] | 22.71 | 0.917 | **0.001** | | | | | | | | | |
| Atmosphere[⁑] | 2.271 | 0.018 | 0.053 | 0.188 | 0.011 | 0.063 | 0.587 | 0.032 | **0.004** | 0.265 | 0.015 | **0.023** |

[†]Mouse identity was controlled for in atmosphere pairwise comparisons.

[‡]Atmosphere was not significant globally for bacterial composition in the vagina when controlling for mouse identity in a subsequent analysis: F = 0.898, $R^2$ = 0.065, p = 0.110.

[⁑]Atmosphere was significant globally for bacterial structure in the vagina when controlling for mouse identity in a subsequent analysis: F = 0.350, $R^2$ = 0.026, p = **0.001**.

*Oral cavity: Oxic vs hypoxic conditions were not significant when controlling for mouse identity in a subsequent analysis: F = 0.681 $R^2$ = 0.039 p = 0.303.

**Supplemental Table 6.** Comparisons of the cultured murine microbiota under (A) anoxic, (B) hypoxic, and (C) oxic conditions.

| **A** | | | | | | Composition | | | | | | | | |
|---|---|---|---|---|---|---|---|---|---|---|---|---|---|---|
| | Anoxic atmosphere | | **Oral cavity n = 11** | | | **Lung n = 9** | | | **Intestine n = 9** | | | **Vagina n = 10** | | |
| | | | **F** | **$R^2$** | **p** | **F** | **$R^2$** | **p** | **F** | **$R^2$** | **p** | **F** | **$R^2$** | **p** |
| | **Oral cavity** | Mouse ID | | | | 1.07 | 0.52 | 0.281 | 1.07 | 0.39 | 0.386 | 0.94 | 0.44 | 0.724 |
| | **Oral cavity** | Body site | | | | 2.08 | 0.10 | **0.004** | 8.46 | 0.31 | **0.001** | 2.83 | 0.13 | **0.001** |
| **Structure** | **Lung** | Mouse ID | 1.14 | 0.51 | 0.281 | | | | 1.06 | 0.46 | 0.358 | 1.05 | 0.56 | 0.264 |
| **Structure** | **Lung** | Body site | 3.04 | 0.14 | **0.006** | | | | 3.94 | 0.19 | **0.001** | 1.42 | 0.08 | **0.041** |
| **Structure** | **Intestine** | Mouse ID | 1.00 | 0.34 | 0.485 | 1.06 | 0.43 | 0.431 | | | | 1.13 | 0.51 | 0.18 |
| **Structure** | **Intestine** | Body site | 10.98 | 0.38 | **0.001** | 5.71 | 0.26 | **0.004** | | | | 3.95 | 0.18 | **0.002** |
| **Structure** | **Vagina** | Mouse ID | 0.90 | 0.40 | 0.588 | 1.28 | 0.55 | 0.208 | 1.14 | 0.46 | 0.326 | | | |
| **Structure** | **Vagina** | Body site | 4.42 | 0.20 | **0.005** | 3.46 | 0.15 | **0.012** | 6.54 | 0.26 | **0.002** | | | |

| **B** | | | | | | Composition* | | | | | | | | |
|---|---|---|---|---|---|---|---|---|---|---|---|---|---|---|
| | Hypoxic atmosphere | | **Oral cavity n = 10** | | | **Lung n = 9** | | | **Intestine n = 10** | | | **Vagina n = 9** | | |
| | | | **F** | **$R^2$** | **p** | **F** | **$R^2$** | **p** | **F** | **$R^2$** | **p** | **F** | **$R^2$** | **p** |
| | **Oral cavity** | Mouse ID | | | | | | | | | | | | |
| | **Oral cavity** | Body site | | | | 2.01 | 0.11 | **0.004** | 4.54 | 0.20 | **0.002** | 2.99 | 0.15 | **0.004** |
| **Structure** | **Lung** | Mouse ID | 1.10 | 0.52 | 0.288 | | | | | | | | | |
| **Structure** | **Lung** | Body site | 3.02 | 0.14 | **0.012** | | | | 2.69 | 0.14 | **0.004** | 1.19 | 0.07 | 0.188 |
| **Structure** | **Intestine** | Mouse ID | 0.68 | 0.343 | 0.928 | 0.82 | 0.47 | 0.841 | | | | | | |
| **Structure** | **Intestine** | Body site | 2.64 | 0.15 | **0.049** | 2.08 | 0.12 | **0.038** | | | | 3.37 | 0.17 | **0.002** |
| **Structure** | **Vagina** | Mouse ID | 0.78 | 0.42 | 0.811 | 0.89 | 0.50 | 0.654 | 1.09 | 0.47 | 0.343 | | | |
| **Structure** | **Vagina** | Body site | 1.51 | 0.09 | 0.185 | 2.96 | 0.17 | **0.025** | 2.88 | 0.14 | **0.019** | | | |

| **C** | | | | | | Composition | | | | | | | | |
|---|---|---|---|---|---|---|---|---|---|---|---|---|---|---|
| | Oxic atmosphere | | **Oral cavity n = 9** | | | **Lung n = 9** | | | **Intestine n = 10** | | | **Vagina n = 10** | | |
| | | | **F** | **$R^2$** | **p** | **F** | **$R^2$** | **p** | **F** | **$R^2$** | **p** | **F** | **$R^2$** | **p** |
| | **Oral cavity** | Mouse ID | | | | 1.09 | 0.51 | 0.243 | 1.00 | 0.42 | 0.490 | 0.99 | 0.44 | 0.498 |
| | **Oral cavity** | Body site | | | | 2.44 | 0.13 | **0.001** | 4.22 | 0.20 | **0.001** | 3.32 | 0.16 | **0.001** |
| **Structure** | **Lung** | Mouse ID | 1.23 | 0.57 | 0.252 | | | | 1.17 | 0.54 | 0.090 | 1.18 | 0.58 | 0.072 |
| **Structure** | **Lung** | Body site | 1.40 | 0.07 | 0.213 | | | | 2.87 | 0.13 | **0.001** | 1.60 | 0.08 | **0.021** |
| **Structure** | **Intestine** | Mouse ID | 0.81 | 0.43 | 0.746 | 1.46 | 0.62 | **0.032**† | | | | 0.99 | 0.43 | 0.537 |
| **Structure** | **Intestine** | Body site | 1.56 | 0.09 | 0.161 | 1.95 | 0.08 | **0.030** | | | | 2.99 | 0.14 | **0.001** |
| **Structure** | **Vagina** | Mouse ID | 0.91 | 0.46 | 0.591 | 2.19 | 0.73 | **0.023**‡ | 0.93 | 0.42 | 0.577 | | | |
| **Structure** | **Vagina** | Body site | 1.53 | 0.09 | 0.218 | 1.20 | 0.04 | 0.323 | 2.51 | 0.13 | **0.042** | | | |

*Because mouse identity had a significant effect on composition under hypoxic conditions, in panel B, all pairwise comparisons of microbiota composition were controlled for mouse identity using the strata function in adonis.

†Lung vs intestine under oxic atmosphere: Body type was significant when controlling for mouse identity in a subsequent analysis: F = 1.769, $R^2$ = 0.094, p = **0.039**.

‡Vagina vs lung under oxic atmosphere: Body type was non-significant when controlling for mouse identity in a subsequent analysis: F = 0.817, $R^2$ = 0.046, p = 0.426.

**Supplemental Table 7.** Prominent amplicon sequence variants (ASVs) shared among cultures from the oral cavity, lung, intestine, and vagina

| Taxonomic classification | ASV | Body site with greatest average relative abundance | Atmosphere with greatest average relative abundance among all body sites |
|---|---|---|---|
| *Rodentibacter* | ASV 2 | Vagina (39.4%) | Oxic (41.4%) |
| *Lactobacillus* | ASV 3 | Lung (26.8%) | Anoxic (20.0%) |
| *Rodentibacter* | ASV 5 | Vagina (31.4%) | Oxic (17.9%) |
| *Staphylococcus* | ASV 17 | Oral cavity (10.4%) | Hypoxic (9.0%) |
| *Rothia nasimurium* | ASV 79 | Vagina (0.93%) | Oxic (1.1%) |

**Supplemental Table 8.** Amplicon sequence variants (ASVs) that were prominent in cultures from only a single body site

| Taxonomic classification | ASV | Body site | Atmosphere with greatest average relative abundance |
|---|---|---|---|
| *Streptococcus danieliae* | ASV 1 | Lung | Anoxic (1.2%) |
| *Staphylococcus* | ASV 8 | Vagina | Hypoxic (5.7%) |
| *Escherichia / Shigella* | ASV 10 | Lung | Anoxic (13.6%) |
| *Streptococcus* | ASV 11 | Oral | Anoxic (2.4%) |
| *Parasutterella* | ASV 20 | Intestine | Anoxic (2.2%) |
| *Bifidobacterium* | ASV 41 | Intestine | Anoxic (1.8%) |
| *Bacteroides acidifaciens* | ASV 53 | Intestine | Anoxic (3.0%) |
| *Acinetobacter* | ASV 60 | Intestine | Oxic (7.1%) |
| *Pseudomonas* | ASV 73 | Intestine | Oxic (8.4%) |
| Actinobacteria | ASV 87 | Lung | Oxic / Hypoxic (6.2%) |
| Streptomycetaceae | ASV 97 | Lung | Oxic (9.9%) |
| Actinobacteria | ASV 106 | Lung | Oxic / Hypoxic (4.8%) |
| *Brevibacillus* | ASV 108 | Lung | Hypoxic (7.1%) |
| *Streptococcus* | ASV 116 | Vagina | Anoxic (1.2%) |
| *Bacillus* | ASV 123 | Intestine | Oxic (4.0%) |
| *Enterorhabdus* | ASV 125 | Lung | Anoxic (6.0%) |
| *Muribacter muris* | ASV 143 | Oral | Oxic (1.1%) |
| *Brevibacillus limnophilus* | ASV 156 | Lung | Hypoxic (3.8%) |
| Coriobacteriaceae UCG-002 | ASV 180 | Intestine | Anoxic (1.1%) |
| *Paenibacillus* | ASV 361 | Intestine | Oxic (1.2%) |

**Supplemental Table 9.** Amplicon sequence variant (ASV) counts from culture and molecular datasets

| | **Culture** | **Molecular** | **Shared** | **% Shared** |
|---|---|---|---|---|
| **Oral cavity** | 177 | 454 | 122 | 24% |
| **Lung** | 152 | 93 | 44 | 22% |
| **Intestine** | 331 | 660 | 262 | 36% |
| **Vagina** | 134 | 510 | 93 | 17% |
| **All body sites** | 411 | 751 | 339 | 41% |

**Supplemental Table 10.** Summary table of the genomes of the two *Rodentibacter* isolates identified in the study alongside type strains and their closest relatives.

| | ***Rodentibacter pneumotropicus*** | | | ***Rodentibacter heylii*** | | |
|---|---|---|---|---|---|---|
| | Type strain | Closest relative | Our strain | Type strain | Closest relative | Our strain |
| **ID** | ATCC 35149 | P441 | ASV2 | ATCC 12555 | G1 | ASV5 |
| **Genome (Mbp)** | 2.43 | 2.4 | 2.45 | 2.72 | 2.61 | 2.51 |
| **G+C content (%)** | 39.87 | 39.81 | 39.9 | 40.2 | 40.19 | 40.13 |
| **Total genes** | 2,297 | 2,296 | 2,384 | 2,656 | 2,520 | 2,474 |
| **Percent of core genome represented** | 100% | 100% | 100% | 100% | 100% | 99.7% (1644 / 1649 genes) |
| **Hypothetical proteins** | 261 | 286 | 308 | 360 | 370 | 285 |

**Supplementary Table 11.** Comparisons of the (A) swab and (B) tissue sample cultured microbiota to technical controls.

| **A** | | | | |
|---|---|---|---|---|
| **Negative swab extraction controls** | **Oral cavity** | | **Vagina** | |
| | **F** | **p** | **F** | **p** |
| **Composition** | 3.569 | **0.001** | 2.482 | **0.001** |
| **Structure** | 17.046 | **0.001** | 8.344 | **0.001** |

| **B** | | | | | | |
|---|---|---|---|---|---|---|
| **Negative extraction controls** | **Lung** | | **Proximal Intestine** | | **Distal Intestine** | |
| | **F** | **p** | **F** | **p** | **F** | **p** |
| **Composition** | 1.521 | **0.018** | 9.723 | **0.001** | 10.021 | **0.001** |
| **Structure** | 2.414 | **0.002** | 7.957 | **0.001** | 8.955 | **0.001** |

**Supplementary Figure 1. Results comparing the influence of atmosphere on the cultured microbiota of the oral cavity of 11 pregnant mice**. In panels A and B, principal coordinates analysis plots indicate bacterial composition (A) and structure (B) of the oral cavity cultured under anoxic, hypoxic, and oxic atmospheres. Ellipses indicate standard deviation for each atmosphere and in panel B, ASVs with the highest relative average abundances were plotted using weighted average scores calculated with the vegan package. The heatmap in panel C includes ASVs with ≥ 1% average relative abundance and samples are clustered by Bray-Curtis similarities within each atmosphere. The cladogram in panel D illustrates LEfSe results of taxa preferentially recovered under one atmosphere over the others. Statistical comparisons were performed using non-parametric MANOVAs with the vegan package in R (v. 4.0.3).

**Supplementary Figure 2. Results comparing the influence of atmosphere on the cultured microbiota of the lung of 10 pregnant mice.** In panels A and B, principal coordinates analysis plots indicate bacterial composition (A) and structure (B) of the lung cultured under anoxic, hypoxic, and oxic atmospheres. The heatmap in panel C includes ASVs with ≥ 1% average relative abundance and samples are clustered by Bray-Curtis similarities. Statistical comparisons were performed using non-parametric MANOVAs with the vegan package in R (v. 4.0.3).

**Supplementary Figure 3. Results comparing the influence of atmosphere on the cultured microbiota of the intestine of 11 pregnant mice.** In panels A and B, principal coordinates analysis plots indicate bacterial composition (A) and structure (B) of the intestine cultured under anoxic, hypoxic, and oxic atmospheres. Ellipses indicate standard deviation for each atmosphere and in panel B, ASVs with the highest relative average abundances were plotted using weighted average scores calculated with the vegan package. The heatmap in panel C includes ASVs with ≥ 1% average relative abundance and samples are clustered by Bray-Curtis similarities within each atmosphere. The cladogram in panel D illustrates LEfSe results of taxa preferentially recovered under one atmosphere over the others. Statistical comparisons were performed using non-parametric MANOVAs with the vegan package in R (v. 4.0.3).

**Supplementary Figure 4. Results comparing the influence of atmosphere on the cultured microbiota of the vagina of 11 pregnant mice.** In panels A and B, principal coordinates analysis plots indicate bacterial composition (A) and structure (B) of the vagina cultured under anoxic, hypoxic, and oxic atmospheres with mouse identity indicated by letters A through K. In panel B, ASVs with the highest relative average abundances were plotted using weighted average scores calculated with the vegan package. The heatmap in panel C includes ASVs with ≥ 1% average relative abundance and samples are clustered by Bray-Curtis similarities. Statistical comparisons were performed using non-parametric MANOVAs with the vegan package in R (v. 4.0.3).

**Supplementary Figure 5. LEfSe results identifying individual ASVs recovered preferentially under one atmosphere over the others for the oral cavity, lung, and intestine.** For each panel Linear discriminate analysis Effect Size (LEfSe) analysis was performed on

ASVs cultured from the indicated body site to identify ASVs preferentially recovered under one atmospheric condition over the others. No ASVs were preferentially recovered under hypoxic or oxic conditions in the lung. No individual ASVs were identified as significant by LEfSe for vaginal samples under any atmosphere.

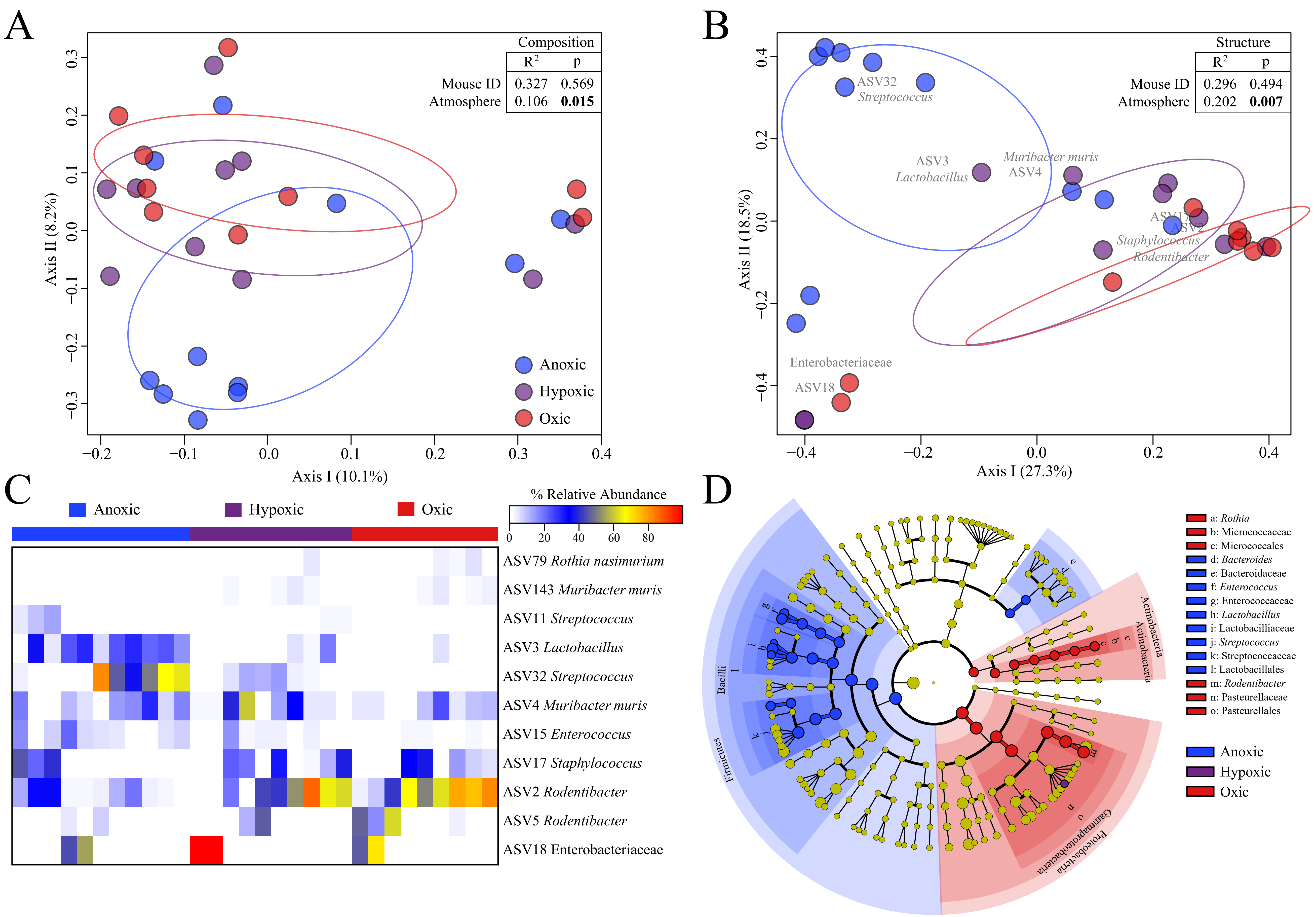

A
Composition
R2 p
Mouse ID 0.327 0.569
Atmosphere 0.106 0.015
Axis II (8.2%)
Axis I (10.1%)
Anoxic
Hypoxic
Oxic
B
Structure
R2 p
Mouse ID 0.296 0.494
Atmosphere 0.202 0.007
ASV32
Streptococcus
ASV3
Lactobacillus
Muribacter muris
ASV4
ASV17
ASV2
Staphylococcus
Rodentibacter
Enterobacteriaceae
ASV18
Axis II (18.5%)
Axis I (27.3%)
C
Anoxic
Hypoxic
Oxic
% Relative Abundance
0 20 40 60 80
ASV79 Rothia nasimurium
ASV143 Muribacter muris
ASV11 Streptococcus
ASV3 Lactobacillus
ASV32 Streptococcus
ASV4 Muribacter muris
ASV15 Enterococcus
ASV17 Staphylococcus
ASV2 Rodentibacter
ASV5 Rodentibacter
ASV18 Enterobacteriaceae
D
a: Rothia
b: Micrococcaceae
c: Micrococcales
d: Bacteroides
e: Bacteroidaceae
f: Enterococcus
g: Enterococcaceae
h: Lactobacillus
i: Lactobacilliaceae
j: Streptococcus
k: Streptococcaceae
l: Lactobacillales
m: Rodentibacter
n: Pasteurellaceae
o: Pasteurellales
Anoxic
Hypoxic
Oxic
Actinobacteria
Bacilli
Firmicutes
Gammaproteobacteria
Proteobacteria

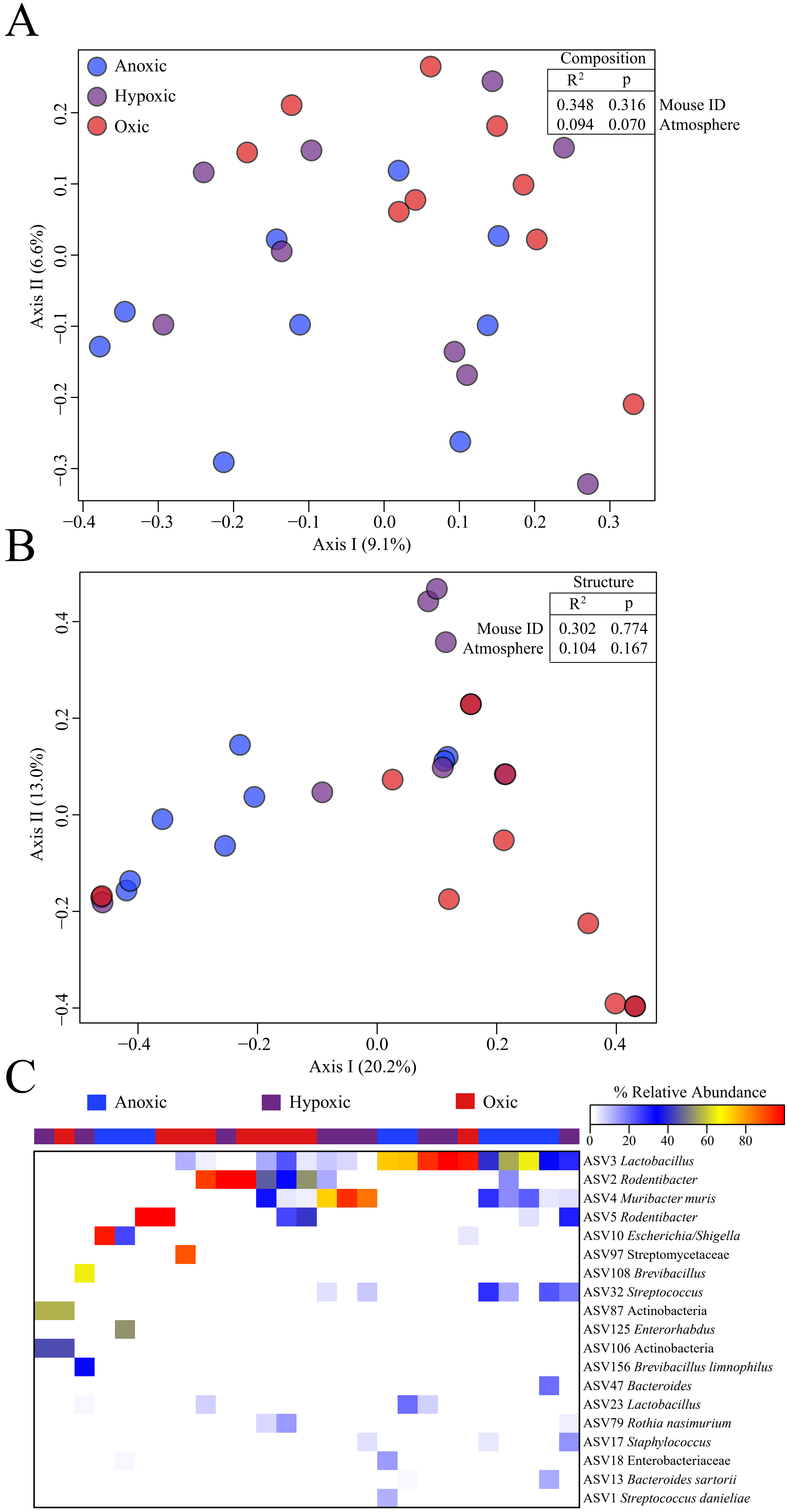

A
Anoxic
Hypoxic
Oxic
Composition
R2 p
0.348 0.316 Mouse ID
0.094 0.070 Atmosphere
Axis II (6.6%)
Axis I (9.1%)
B
Structure
R2 p
Mouse ID 0.302 0.774
Atmosphere 0.104 0.167
Axis II (13.0%)
Axis I (20.2%)
C
Anoxic
Hypoxic
Oxic
% Relative Abundance
0 20 40 60 80
ASV3 Lactobacillus
ASV2 Rodentibacter
ASV4 Muribacter muris
ASV5 Rodentibacter
ASV10 Escherichia/Shigella
ASV97 Streptomycetaceae
ASV108 Brevibacillus
ASV32 Streptococcus
ASV87 Actinobacteria
ASV125 Enterorhabdus
ASV106 Actinobacteria
ASV156 Brevibacillus limnophilus
ASV47 Bacteroides
ASV23 Lactobacillus
ASV79 Rothia nasimurium
ASV17 Staphylococcus
ASV18 Enterobacteriaceae
ASV13 Bacteroides sartorii
ASV1 Streptococcus danieliae

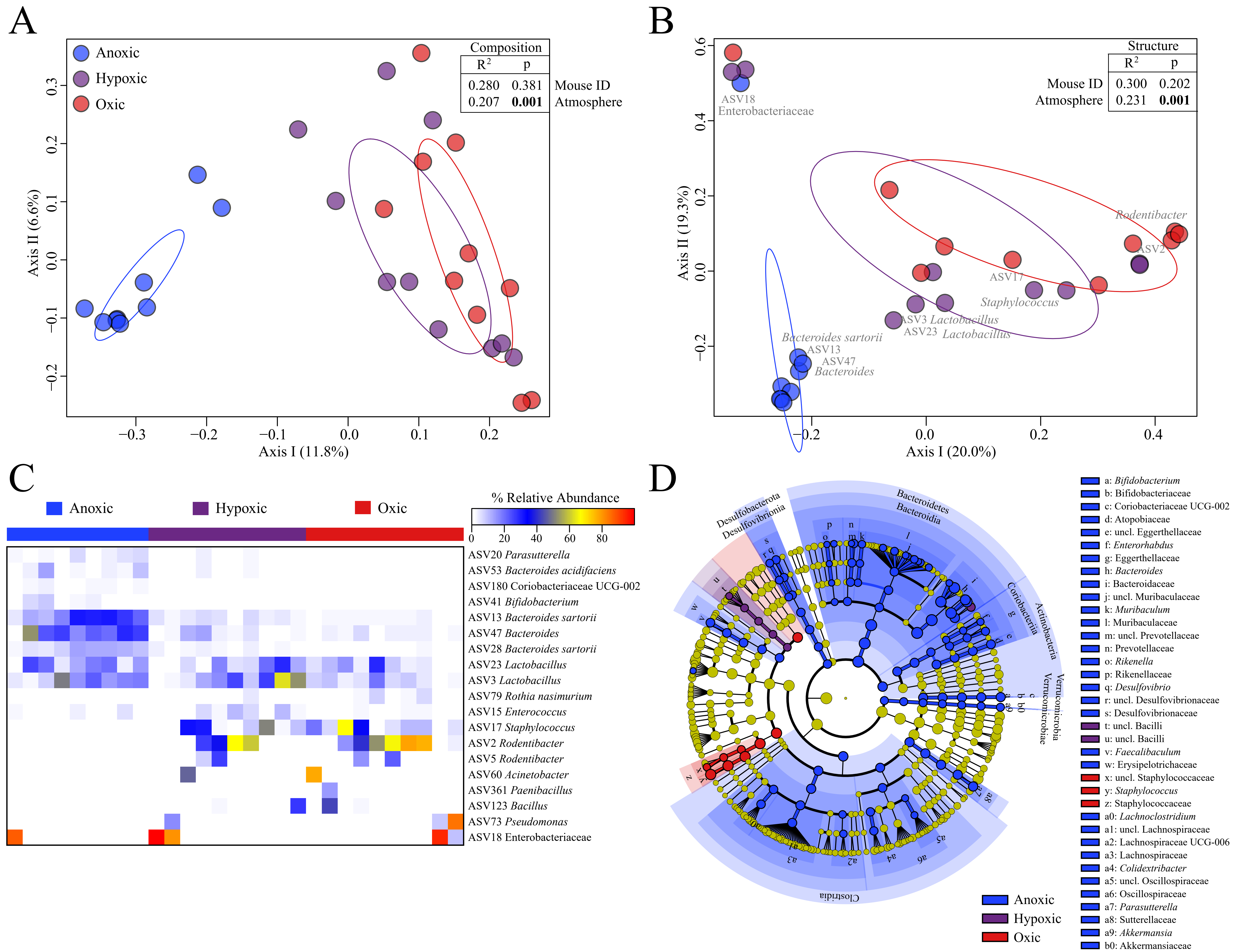
A
Anoxic
Hypoxic
Oxic
Composition
R2 p
0.280 0.381 Mouse ID
0.207 0.001 Atmosphere
Axis II (6.6%)
Axis I (11.8%)
B
Structure
R2 p
Mouse ID 0.300 0.202
Atmosphere 0.231 0.001
ASV18
Enterobacteriaceae
Rodentibacter
ASV2
ASV17
Staphylococcus
ASV3 Lactobacillus
ASV23 Lactobacillus
Bacteroides sartorii
ASV13
ASV47
Bacteroides
Axis II (19.3%)
Axis I (20.0%)
C
Anoxic
Hypoxic
Oxic
% Relative Abundance
0 20 40 60 80
ASV20 Parasutterella
ASV53 Bacteroides acidifaciens
ASV180 Coriobacteriaceae UCG-002
ASV41 Bifidobacterium
ASV13 Bacteroides sartorii
ASV47 Bacteroides
ASV28 Bacteroides sartorii
ASV23 Lactobacillus
ASV3 Lactobacillus
ASV79 Rothia nasimurium
ASV15 Enterococcus
ASV17 Staphylococcus
ASV2 Rodentibacter
ASV5 Rodentibacter
ASV60 Acinetobacter
ASV361 Paenibacillus
ASV123 Bacillus
ASV73 Pseudomonas
ASV18 Enterobacteriaceae
D
Desulfobacterota
Desulfovibrionia
Bacteroidetes
Bacteroidia
Coriobacteriia
Actinobacteria
Verrucomicrobia
Verrucomicrobiae
Clostridia
a: Bifidobacterium
b: Bifidobacteriaceae
c: Coriobacteriaceae UCG-002
d: Atopobiaceae
e: uncl. Eggerthellaceae
f: Enterorhabdus
g: Eggerthellaceae
h: Bacteroides
i: Bacteroidaceae
j: uncl. Muribaculaceae
k: Muribaculum
l: Muribaculaceae
m: uncl. Prevotellaceae
n: Prevotellaceae
o: Rikenella
p: Rikenellaceae
q: Desulfovibrio
r: uncl. Desulfovibrionaceae
s: Desulfovibrionaceae
t: uncl. Bacilli
u: uncl. Bacilli
v: Faecalibaculum
w: Erysipelotrichaceae
x: uncl. Staphylococcaceae
y: Staphylococcus
z: Staphylococcaceae
a0: Lachnoclostridium
a1: uncl. Lachnospiraceae
a2: Lachnospiraceae UCG-006
a3: Lachnospiraceae
a4: Colidextribacter
a5: uncl. Oscillospiraceae
a6: Oscillospiraceae
a7: Parasutterella
a8: Sutterellaceae
a9: Akkermansia
b0: Akkermansiaceae
Anoxic
Hypoxic
Oxic

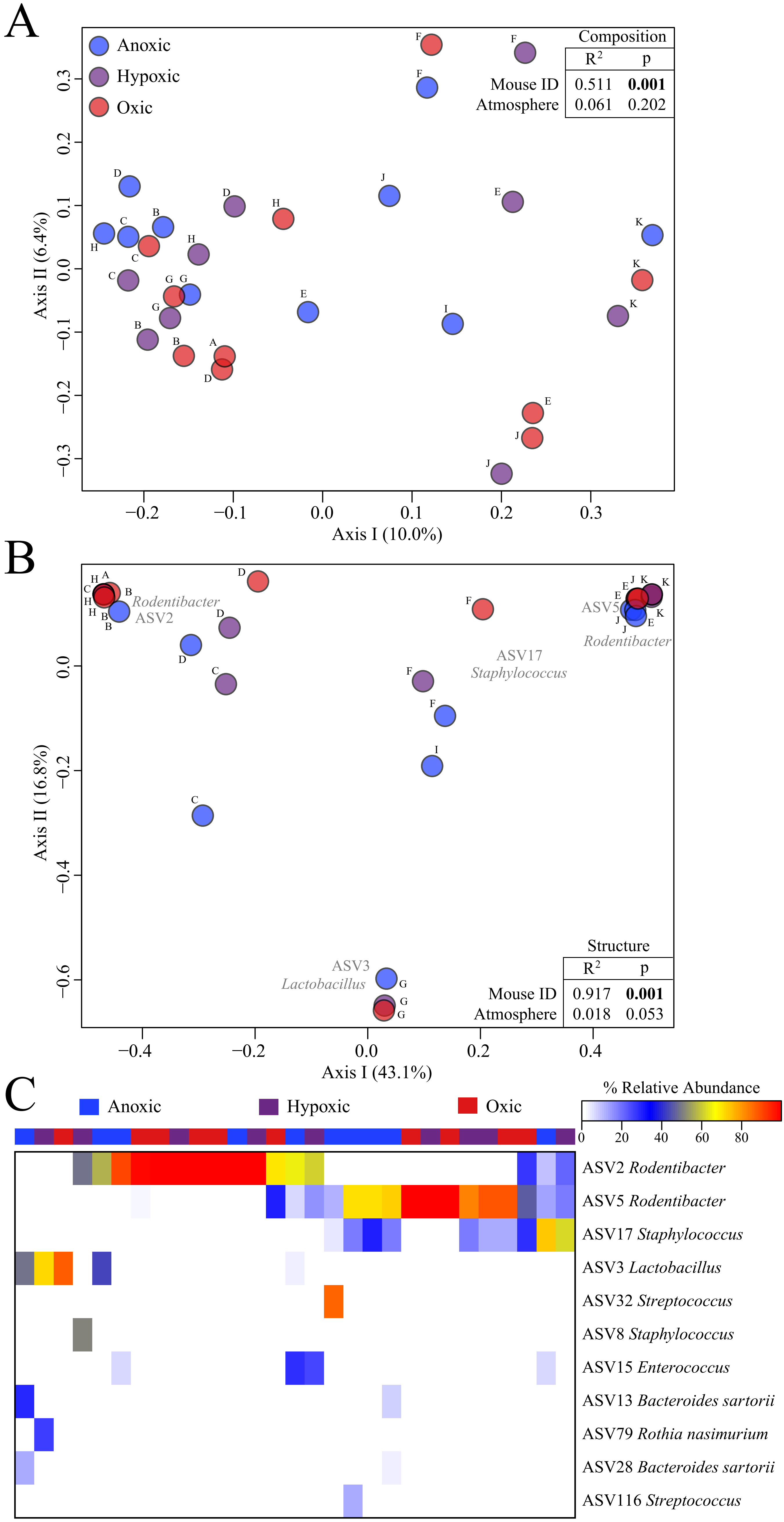

A
Anoxic
Hypoxic
Oxic
Composition
R2 p
Mouse ID 0.511 0.001
Atmosphere 0.061 0.202
Axis I (10.0%)
Axis II (6.4%)
B
Rodentibacter ASV2
ASV5 Rodentibacter
ASV17 Staphylococcus
ASV3 Lactobacillus
Structure
R2 p
Mouse ID 0.917 0.001
Atmosphere 0.018 0.053
Axis I (43.1%)
Axis II (16.8%)
C
Anoxic
Hypoxic
Oxic
% Relative Abundance
0 20 40 60 80
ASV2 Rodentibacter
ASV5 Rodentibacter
ASV17 Staphylococcus
ASV3 Lactobacillus
ASV32 Streptococcus
ASV8 Staphylococcus
ASV15 Enterococcus
ASV13 Bacteroides sartorii
ASV79 Rothia nasimurium
ASV28 Bacteroides sartorii
ASV116 Streptococcus

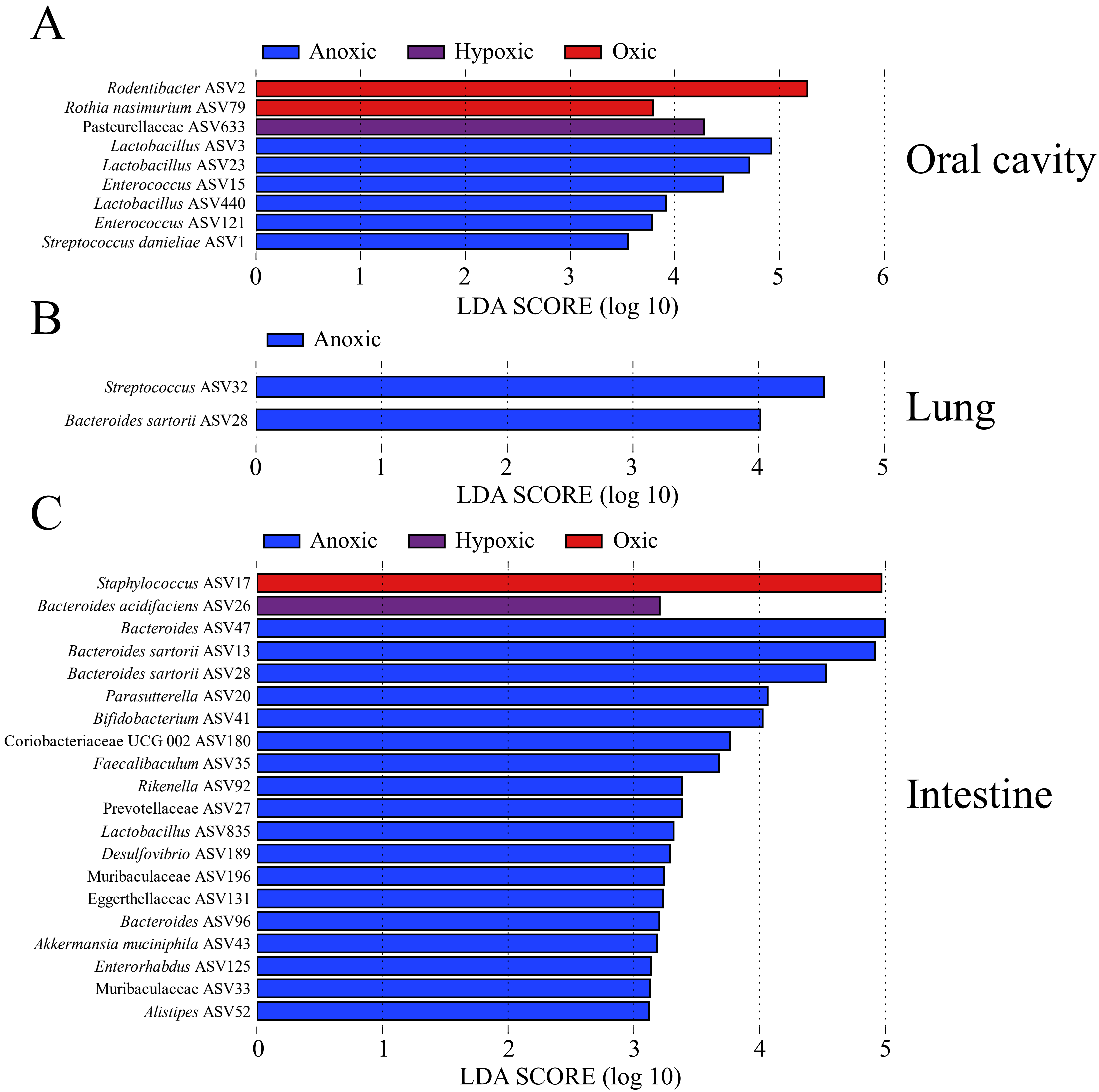
A
Anoxic
Hypoxic
Oxic
Rodentibacter ASV2
Rothia nasimurium ASV79
Pasteurellaceae ASV633
Lactobacillus ASV3
Lactobacillus ASV23
Enterococcus ASV15
Lactobacillus ASV440
Enterococcus ASV121
Streptococcus danieliae ASV1
0
1
2
3
4
5
6
LDA SCORE (log 10)
Oral cavity
B
Anoxic
Streptococcus ASV32
Bacteroides sartorii ASV28
0
1
2
3
4
5
LDA SCORE (log 10)
Lung
C
Anoxic
Hypoxic
Oxic
Staphylococcus ASV17
Bacteroides acidifaciens ASV26
Bacteroides ASV47
Bacteroides sartorii ASV13
Bacteroides sartorii ASV28
Parasutterella ASV20
Bifidobacterium ASV41
Coriobacteriaceae UCG 002 ASV180
Faecalibaculum ASV35
Rikenella ASV92
Prevotellaceae ASV27
Lactobacillus ASV835
Desulfovibrio ASV189
Muribaculaceae ASV196
Eggerthellaceae ASV131
Bacteroides ASV96
Akkermansia muciniphila ASV43
Enterorhabdus ASV125
Muribaculaceae ASV33
Alistipes ASV52
0
1
2
3
4
5
LDA SCORE (log 10)
Intestine

### References

1. Abusleme L, Hong B-Y, Hoare A, Konkel JE, Diaz PI, Moutsopoulos NM. 2017. Oral Microbiome Characterization in Murine Models. Bio-protocol 7:e2655.
2. Hernández-Arriaga A, Baumann A, Witte OW, Frahm C, Bergheim I, Camarinha-Silva A. 2019. Changes in Oral Microbial Ecology of C57BL/6 Mice at Different Ages Associated with Sampling Methodology. Microorganisms 7:283.
3. Abusleme L, O'Gorman H, Dutzan N, Greenwell-Wild T, Moutsopoulos NM. 2020. Establishment and Stability of the Murine Oral Microbiome. Journal of Dental Research 99:721-729.
4. Jašarević E, Hill EM, Kane PJ, Rutt L, Gyles T, Folts L, Rock KD, Howard CD, Morrison KE, Ravel J, Bale TL. 2021. The composition of human vaginal microbiota transferred at birth affects offspring health in a mouse model. Nature Communications 12:6289.
5. Barfod KK, Roggenbuck M, Hansen LH, Schjørring S, Larsen ST, Sørensen SJ, Krogfelt KA. 2013. The murine lung microbiome in relation to the intestinal and vaginal bacterial communities. BMC microbiology 13:303-303.
6. Poroyko V, Meng F, Meliton A, Afonyushkin T, Ulanov A, Semenyuk E, Latif O, Tesic V, Birukova AA, Birukov KG. 2015. Alterations of lung microbiota in a mouse model of LPS-induced lung injury. American journal of physiology Lung cellular and molecular physiology 309:L76-L83.
7. Singh N, Vats A, Sharma A, Arora A, Kumar A. 2017. The development of lower respiratory tract microbiome in mice. Microbiome 5:61-61.
8. Dickson RP, Erb-Downward JR, Falkowski NR, Hunter EM, Ashley SL, Huffnagle GB. 2018. The Lung Microbiota of Healthy Mice Are Highly Variable, Cluster by Environment, and Reflect Variation in Baseline Lung Innate Immunity. American journal of respiratory and critical care medicine 198:497-508.
9. Kostric M, Milger K, Krauss-Etschmann S, Engel M, Vestergaard G, Schloter M, Schöler A. 2018. Development of a Stable Lung Microbiome in Healthy Neonatal Mice. Microbial Ecology 75:529-542.
10. Ashley SL, Sjoding MW, Popova AP, Cui TX, Hoostal MJ, Schmidt TM, Branton WR, Dieterle MG, Falkowski NR, Baker JM, Hinkle KJ, Konopka KE, Erb-Downward JR, Huffnagle GB, Dickson RP. 2020. Lung and gut microbiota are altered by hyperoxia and contribute to oxygen-induced lung injury in mice. Science Translational Medicine 12:eaau9959.
11. Li J, Hu Y, Liu L, Wang Q, Zeng J, Chen C. 2020. PM2.5 exposure perturbs lung microbiome and its metabolic profile in mice. Sci Total Environ 721:137432.
12. Baker JM, Hinkle KJ, McDonald RA, Brown CA, Falkowski NR, Huffnagle GB, Dickson RP. 2021. Whole lung tissue is the preferred sampling method for amplicon-based characterization of murine lung microbiota. Microbiome 9:99.
13. Lipinski JH, Falkowski NR, Huffnagle GB, Erb-Downward JR, Dickson RP, Moore BB, O'Dwyer DN. 2021. Toll-like receptors, environmental caging, and lung dysbiosis. American Journal of Physiology-Lung Cellular and Molecular Physiology 321:L404-L415.
14. Oh JE, Kim B-C, Chang D-H, Kwon M, Lee SY, Kang D, Kim JY, Hwang I, Yu J-W, Nakae S, Lee HK. 2016. Dysbiosis-induced IL-33 contributes to impaired antiviral

immunity in the genital mucosa. Proceedings of the National Academy of Sciences 113:E762-E771.

15. Vrbanac A, Riestra AM, Coady A, Knight R, Nizet V, Patras KA. 2018. The murine vaginal microbiota and its perturbation by the human pathogen group B Streptococcus. BMC Microbiology 18:197.
16. Karpinets TV, Solley TN, Mikkelson MD, Dorta-Estremera S, Nookala SS, Medrano AYD, Petrosino JF, Mezzari MP, Zhang J, Futreal PA, Sastry KJ, Colbert LE, Klopp A. 2020. Effect of Antibiotics on Gut and Vaginal Microbiomes Associated with Cervical Cancer Development in Mice. Cancer Prevention Research 13:997-1006.
17. Hernández-Quiroz F, Murugesan S, Velazquez-Martínez C, Villalobos-Flores LE, Maya-Lucas O, Piña-Escobedo A, García-González I, Ocadiz-Delgado R, Lambert PF, Gariglio P, García-Mena J. 2021. The vaginal and fecal microbiota of a murine cervical carcinoma model under synergistic effect of 17β-Estradiol and E7 oncogene expression. Microbial Pathogenesis 152:104763.
18. Mejia ME, Ottinger S, Vrbanac A, Babu P, Zulk JJ, Moorshead D, Bode L, Nizet V, Patras KA, Fey PD. 2022. Human Milk Oligosaccharides Reduce Murine Group B *Streptococcus* Vaginal Colonization with Minimal Impact on the Vaginal Microbiota. mSphere 7:e00885-21.
